# Topological reprogramming transforms an integral membrane oligosaccharyltransferase into a water-soluble glycosylation catalyst

**DOI:** 10.64898/2026.01.30.702934

**Authors:** Yong Hyun Kwon, Ljubica Mihaljević, Keehun Kim, David E. Kim, Thomas C. Donahue, Erik J. Bidstrup, Chandra K. Bandi, Belen Sotomayor, Sophia W. Hulbert, Kathryn A. Myers, Anru Tian, Mariah Culpepper, Dario Mizrachi, Thapakorn Jaroentomeechai, Henrik Clausen, Michael C. Jewett, David Baker, Matthew P. DeLisa

## Abstract

Glycosyltransferases (GTs) catalyze the formation of new glycosidic bonds and thus are vital for synthesizing nature’s vast repertoire of glycans and glycoconjugates and for engineering glycan-related medicines and materials. However, obtaining detailed structural and functional insights for the >750,000 known GTs is limited by difficulties associated with their efficient recombinant expression. Members of the GT-C fold, in particular, pose the most significant expression challenges due to the integration and folding requirements of their multiple membrane-spanning regions. Here, we address this challenge by engineering water-soluble variants of an archetypal GT-C fold enzyme, namely the oligosaccharyltransferase PglB from *Campylobacter jejuni* (*Cj*PglB), which possesses 13 hydrophobic transmembrane helices. To render *Cj*PglB water-soluble, we leveraged two advanced protein engineering methods: one that is universal called SIMPLEx (<u>s</u>olubilization of <u>IMP</u>s with high levels of <u>ex</u>pression) and the other that is custom tailored called WRAPs (<u>w</u>ater-soluble <u>R</u>Fdiffused <u>a</u>mphipathic <u>p</u>roteins). Each approach was able to transform *Cj*PglB into a water-soluble enzyme that could be readily expressed in the cytoplasm of *Escherichia coli* cells at yields in the 3–6 mg/L range. Importantly, solubilization was achieved without the need for detergents and with retention of catalytic function. Collectively, our findings demonstrate that both SIMPLEx and WRAPs are promising platforms for advancing the molecular characterization of even the most structurally complex GTs, while also enabling broader GT-mediated glycosylation capabilities within synthetic glycobiology applications.

## Introduction

Glycosyltransferases (GTs) are ubiquitous enzymes that catalyze the transfer of sugar moieties from activated donor molecules to specific acceptor substrates, resulting in the formation of glycosidic bonds ^1^. The diverse and extensive collection of GTs across all domains of life includes more than 750,000 known sequences ^2, 3^, which encompass three main structural superfamilies–GT-A, GT-B, and GT-C–that are distinguished by unique folds and specific modes of substrate interaction ^1, 4, 5^. These enzymes use different substrates including nucleotide sugars (Leloir enzymes) and lipid-linked sugars (non-Leloir enzymes), and act on a range of acceptors including sugars, proteins, lipids, and small molecules. This broad enzymatic capability represents a vital toolkit for glycosylation processes in nature ^1–5^ and biotechnology ^6–13^.

Despite the vast repertoire, relatively few GTs have been characterized at the molecular level through structural and kinetic studies, leaving much of their molecular diversity poorly understood and restricting the full scope of their biotechnological potential. This gap has been ascribed to multiple challenges; however, one of the most significant, given its crucial role early in the characterization workflow, is the recombinant expression of GTs as functional proteins ^4, 5, 14–16^. One reason recombinant expression is so difficult stems from the fact that many GTs are membrane-associated, either via peripheral or single-pass anchoring in the case of GT-A and GT-B folds or multi-transmembrane helices in the case of GT-C folds. Consequently, many GTs are intrinsically hydrophobic and depend on membrane environments, leading to misfolding and aggregation following recombinant expression and poor *in vitro* solubility and stability following purification. While GT expression in mammalian, insect, and yeast cells has met some success, including two larger-scale expression campaigns ^15, 17^, there remains a need for a broadly effective expression platform that can be readily generalized for rapid, high-yield production of functional GTs from all three structural superfamilies.

To this end, we recently described a strategy for efficient production of structurally diverse GTs that leverages the biosynthetic capacity and versatility of *Escherichia coli* bacteria, the workhorse of modern molecular biology ^18^. Our approach centered around SIMPLEx (<u>s</u>olubilization of <u>IMP</u>s with high levels of <u>ex</u>pression), a universal protein engineering method for topologically converting peripheral and integral membrane proteins (IMPs) into water-soluble variants ^19^. To adapt GTs for SIMPLEx, chimeras were created by genetically fusing candidate glycoenzymes with a decoy protein at their N-termini that prevented membrane insertion and an amphipathic protein at their C-termini that effectively shielded hydrophobic surfaces from the aqueous environment ^16^. Our approach promoted soluble expression of nearly 100 GTs, including many of human origin, directly in the *E. coli* cytoplasm. Importantly, water-soluble GTs were all produced at high yields (5–10 mg/L) without the need for added detergents and with intact catalytic function.

In our previous work, we focused exclusively on enzymes from the GT-A and GT-B superfamilies, which use nucleotide sugars as donor substrates and only contain between 0 and 1 transmembrane domains (TMDs) on average ^16^. In stark contrast, GT-C fold enzymes use lipid phosphate-activated donor sugar substrates and have between 8 and 13 TMDs with an active site located within a long-loop region ^1, 20^, making them more topologically complex than other GT folds. Because of their complicated fold architectures, detailed experimental information about the structure and function of integral membrane GT-C fold enzymes is still sparse ^21^.

Therefore, in this work, we investigated whether the SIMPLEx-mediated solubilization approach could be used to generate water-soluble variants of GT-C superfamily members. As proof-of-concept, we focused on the PglB enzyme from *Campylobacter jejuni* (*Cj*PglB), a bacterial single-subunit oligosaccharyltransferase (OST) that serves as an archetype of the GT-C fold, consisting of 13 TMDs and a conserved C-terminal globular domain that harbors the active site and the hallmark WWDYG motif critical for catalytic function ^22^. *Cj*PglB and its structural homologues (e.g., archaeal AglB, eukaryotic STT3 catalytic subunit) mediate the key step of asparagine-linked (*N*-linked) glycosylation by transferring preassembled glycans from polyisoprenol-linked donors onto specific asparagine residues in acceptor proteins ^23,24^. Following several modifications to the original SIMPLEx design, we identified an optimized architecture that conferred water solubility to *Cj*PglB in the absence of detergents, with final yields exceeding 5 mg/L. Importantly, the catalytic activity of SIMPLEx-solubilized *Cj*PglB rivaled that of detergent-solubilized wild-type (wt) *Cj*PglB, with both enzymes exhibiting ∼100% glycosylation efficiency using a model acceptor protein substrate and exhibiting comparable Michaelis-Menten kinetics using a peptide substrate. In parallel, we also investigated an alternative deep learning-based design approach called WRAPs (<u>w</u>ater-soluble <u>R</u>Fdiffused <u>a</u>mphipathic <u>p</u>roteins) ^25^ for custom solubilization of *Cj*PglB. Similar to SIMPLEx, the best WRAP design was also able to render *Cj*PglB water-soluble without the need for detergents and with retention of function, indicating that there are multiple solutions to the water-soluble protein design problem. Taken together, our results demonstrate two promising platforms–SIMPLEx and WRAPs–for unlocking structure-function relationships of GT-C fold enzymes as well as expanding glycosylation capabilities in synthetic glycobiology.

## Results

### Engineering water-soluble *Cj*PglB variants using SIMPLEx technology

We initially focused our attention on *Cj*PglB, a GT-C fold family member consisting of 13 TMDs (**Supplementary Fig. 1a**). To render *Cj*PglB water soluble, we designed a tripartite chimera based on the original SIMPLEx architecture ^16, 19^ in which the N-terminus of *Cj*PglB was genetically fused to a water-soluble “decoy” protein, namely ΔspMBP (*E. coli* maltose-binding protein lacking its N-terminal signal peptide), and the C-terminus was fused to an amphipathic “shield” protein, namely ApoAI* (truncated human apolipoprotein A1 lacking its 43-residue globular N-terminal domain) (**Fig. 1a**). However, following expression in *E. coli* BL21(DE3) cells, the ΔspMBP-*Cj*PglB-ApoAI* chimera (hereafter Sx-*Cj*PglB.v1) accumulated almost entirely in the detergent-solubilized membrane fraction with virtually no detectable accumulation in the soluble fraction (**Supplementary Fig. 1b**). In fact, the expression pattern for Sx-*Cj*PglB.v1 was similar to that observed for wild-type (wt) *Cj*PglB without any fusion partners (**Fig. 1b**), indicating that the original SIMPLEx design was incapable of solubilizing the OST. These results were in stark contrast to what was seen for SIMPLEx-reformatted variants of human ST6Gal1 and FUT8 from the GT-A and GT-B superfamilies, respectively, which both accumulated predominantly in the soluble fraction (**Supplementary Fig. 1b**), as we had seen previously ^16^.

**Figure 1.**
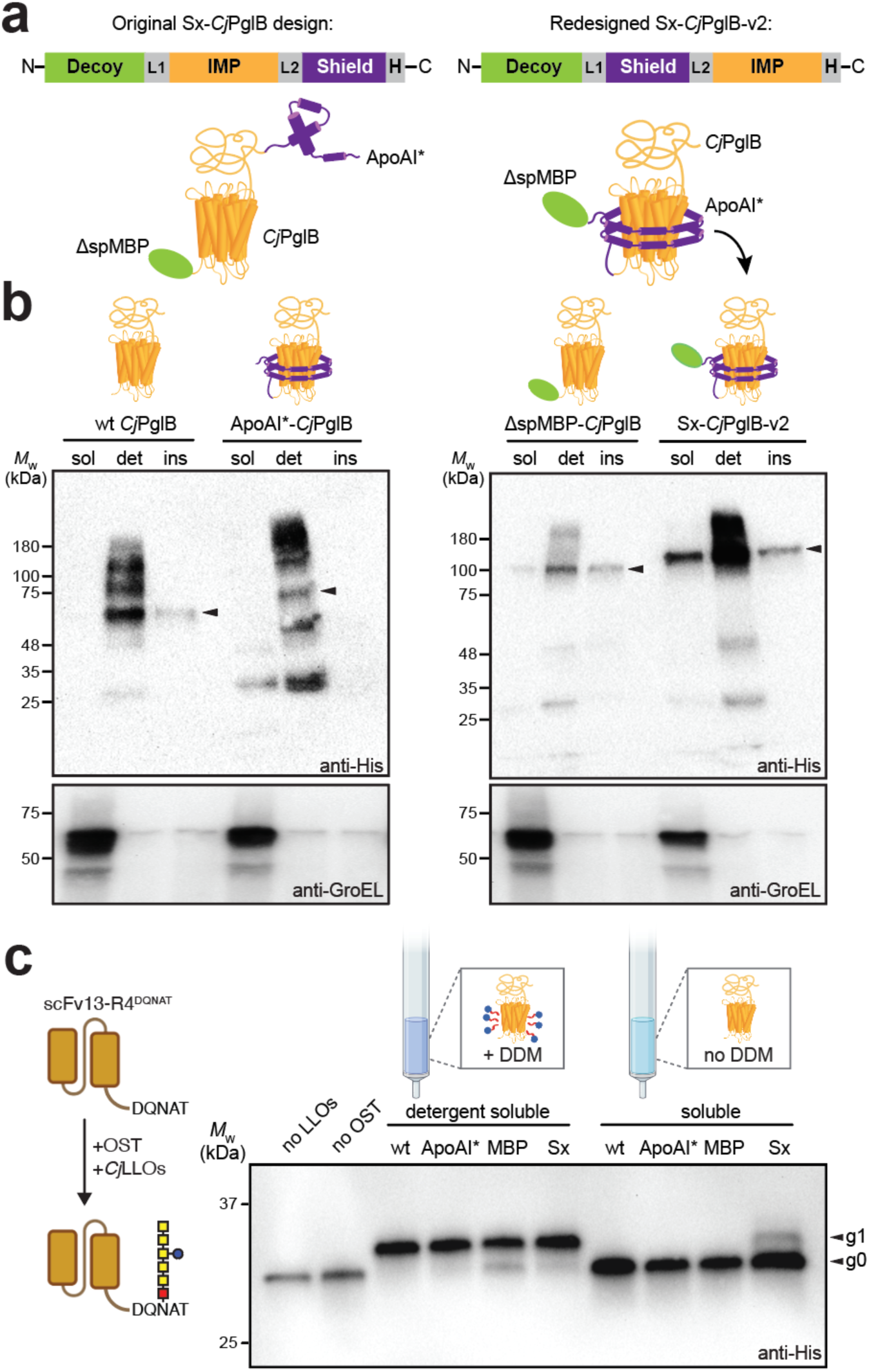
Redesigned SIMPLEx architecture enables solubilization of *Cj*PglB. (a) Schematic of original and redesigned SIMPLEx architectures (see text for details). L1 and L2: two-residue linkers created by SacI and NcoI restriction sites, respectively; H: 10xHis tag. (b) Immunoblot analysis of soluble (sol), detergent solubilized (det), and insoluble (ins) fractions prepared from *E. coli* BL21(DE3) cells expressing each construct. An equivalent amount of total protein was loaded in each lane. Blots were probed with anti-His antibody to detect each of the expressed constructs and anti-GroEL antibody to confirm quality of fractionations and equivalent loading of soluble fractions. Blot on left required much longer exposure time than blot on right. (c) Immunoblot analysis of IVG reactions in which different *Cj*PglB constructs were purified from soluble (sol) or detergent solubilized (det) fractions of BL21(DE3) cells and incubated with organic solvent-extracted *Cj*LLOs and purified scFv13-R4^DQNAT^ acceptor protein. IVG reactions performed without *Cj*LLOs or *Cj*PglB served as negative controls. Blots were probed with anti-His to detect the acceptor protein. Black arrows denote aglycosylated (g0) and singly glycosylated (g1) forms of scFv13-R4^DQNAT^. For all immunoblots, molecular weight (*M*_W_) markers are indicated at left and results are representative of biological replicates (*n* = 3).

We suspected that the lack of soluble expression was due to the complex topology of *Cj*PglB. Specifically, *Cj*PglB possesses a globular periplasmic domain that comprises the C-terminal half of the enzyme and accounts for ∼40% of its mass. In the original Sx-*Cj*PglB.v1 design, this large, extramembranous domain sits between the solubilizing ApoAI* domain and the 13 hydrophobic TMDs of *Cj*PglB. Such an arrangement is likely suboptimal for promoting intimate contact between ApoAI* and the TMDs of the target IMP, which is a key feature of SIMPLEx-mediated solubilization ^19^. Therefore, we redesigned the SIMPLEx architecture by moving the ApoAI* domain to the N-terminal side of *Cj*PglB (**Fig. 1a**), which we hypothesized would provide ApoAI* with less obstructed access to the TMDs of *Cj*PglB. When the newly redesigned ΔspMBP-ApoAI*-*Cj*PglB construct (hereafter Sx-*Cj*PglB.v2) was expressed in BL21(DE3) cells, we readily detected soluble expression in the cytoplasm without the need for added detergents (**Fig. 1b**), indicating that *Cj*PglB had been converted into a water-soluble conformation. Omission of the decoy protein resulted in no detectable accumulation in the soluble fraction, with the decoy-less ApoAI*-*Cj*PglB construct partitioning almost exclusively to the detergent-solubilized membrane fraction in a manner that was identical to unfused wt *Cj*PglB. It is noteworthy that expression of these two constructs was significantly lower than Sx-*Cj*PglB.v2, requiring much longer exposure of immunoblots to generate equivalent band intensities. Omission of ApoAI* resulted in modest levels of expression in all three fractions with just a small portion accumulating in the soluble fraction that likely corresponded to soluble aggregates of *Cj*PglB, which was seen for a previously tested decoy-IMP fusion ^19^. Overall, our results agree with previous findings ^16, 19^ that both the decoy protein and amphipathic shield protein are equally important for effective IMP solubilization. In addition to solubility, we also investigated purification yield given the strong expression of Sx-*Cj*PglB.v2 relative to the other constructs. Following expression and purification from BL21(DE3) cells, the yield of Sx-*Cj*PglB.v2 reached 5.1 mg/L (**Supplementary Fig. 2**), which aligned closely with the 5–10 mg/L yield values reported previously for other SIMPLEx-reformatted IMPs ^16, 19^. Conversely, the yield of wt *Cj*PglB was only 0.5 mg/L, revealing that the SIMPLEx strategy not only promoted solubilization of *Cj*PglB but also increased its purification yield by an order of magnitude compared to the unfused enzyme.

We next determined whether the water-soluble Sx-*Cj*PglB.v2 construct retained enzymatic activity. To this end, we implemented an established *in vitro* glycosylation (IVG) assay ^26^, which involved purifying each *Cj*PglB construct from the different cell fractions and subsequently combining the purified enzymes with their requisite glycosylation substrates. Specifically, lipid-linked oligosaccharides (LLOs) bearing the *C. jejuni* heptasaccharide *N-*glycan (*Cj*LLOs) with a mixture of diNAcBac and GlcNAc as the reducing-end sugar (i.e., GalNAc_5_(Glc)diNAcBac/GlcNAc, where diNAcBac = bacillosamine) were organic solvent-extracted from glycoengineered *E. coli* and used as the donor substrate. A purified single-chain Fv antibody fragment harboring two mutations to remove putative internal glycosylation sites and a DQNAT glycosylation tag and 6xHis at its C-terminus (scFv13-R4(N34L, N77L)^DQNAT-His^^6^; hereafter scFv13-R4^DQNAT^) served as the acceptor protein substrate. To evaluate the glycosylation status of scFv13-R4^DQNAT^ in IVG reactions, immunoblot analysis was performed using an anti-His antibody to detect the formation of aglycosylated (g0) and mono-glycosylated (g1) forms of the scFv13-R4^DQNAT^ acceptor protein. According to this analysis, productive glycosylation was achieved with Sx-*Cj*PglB.v2 that was purified from the soluble and detergent-solubilized membrane fractions (**Fig. 1c**). Importantly, because proteins purified from the soluble portion of the lysate were not supplemented with detergent, the glycosylation activity observed for this fraction was attributed to spontaneously soluble Sx-*Cj*PglB.v2 (i.e., SIMPLEx-reformatted *Cj*PglB that could dissolve in water without requiring additional treatments to aid dissolution). In stark contrast, wt *Cj*PglB, ApoAI*-*Cj*PglB, and ΔspMBP-*Cj*PglB only exhibited activity when purified from the detergent-solubilized membrane fraction (**Fig. 1c**), consistent with their significant accumulation in this fraction. Interestingly, the ΔspMBP-*Cj*PglB construct purified from the soluble fraction showed little to no glycosylation activity, despite its modest accumulation in this fraction, providing further evidence that this construct was in a partially soluble but misfolded state. Collectively, these data establish that our SIMPLEx methodology is useful for solubilizing a member of the GT-C superfamily, *Cj*PglB, while preserving its sequence, fold, and function.

### Modularity of SIMPLEx enables fine-tuning of solubility and catalytic activity

Although the Sx-*Cj*PglB.v2 construct purified from the soluble fraction was active, its glycosylation efficiency (measured as the intensity of the g1 band divided by the sum of the intensities of the g0 and g1 bands in the immunoblot in **Fig. 1c**) was considerably lower than unfused wt *Cj*PglB purified from the detergent-solubilized membrane fraction (22% vs 100% efficient, respectively) under the conditions tested. Therefore, we sought to enhance the efficiency of the water-soluble *Cj*PglB construct by leveraging the modularity of the SIMPLEx architecture. Specifically, we generated a panel of variants in which the size of the decoy domain was decreased while the size of the amphipathic shield domain was increased (**Fig. 2a**). In the case of the decoy, we replaced the ∼42-kDa ΔspMBP domain with smaller hydrophilic sequences, namely small ubiquitin-like modifier protein (SUMO; ∼11 kDa) or a triply repeated FLAG epitope (DYKDHDGDYKDHDIDYKDDDDK; ∼1 kDa). We chose these domains because their high intrinsic solubility and compact size are well known to enhance expression and purification of their fusion partners without impacting activity ^27, 28^. In the case of the shield, we incrementally increased the size of the ApoAI* domain, which naturally contains 10 amphipathic helices (10H) ^29^. Specifically, head-to-tail fusion of ApoAI* with the C-terminal half of ApoAI* containing helices 6–10 resulted in a 15-helix (15H) design while fusing an additional full-length copy of ApoAI* containing helices 1–10 resulted in a 20-helix (20H) design. The rationale for lengthening this domain was based on our previous low-resolution structural analysis, which showed that ApoAI* tended to wrap around the transmembrane helices of its IMP fusion partner ^19^, consistent with the “belt”-like conformation it adopts with phospholipids ^30^. To shield the 13 TMDs of *Cj*PglB, which present considerably more hydrophobic surface area than any of the previously solubilized IMPs (which possessed between 3–7 TMDs), we speculated that a longer belt might be needed.

**Figure 2.**
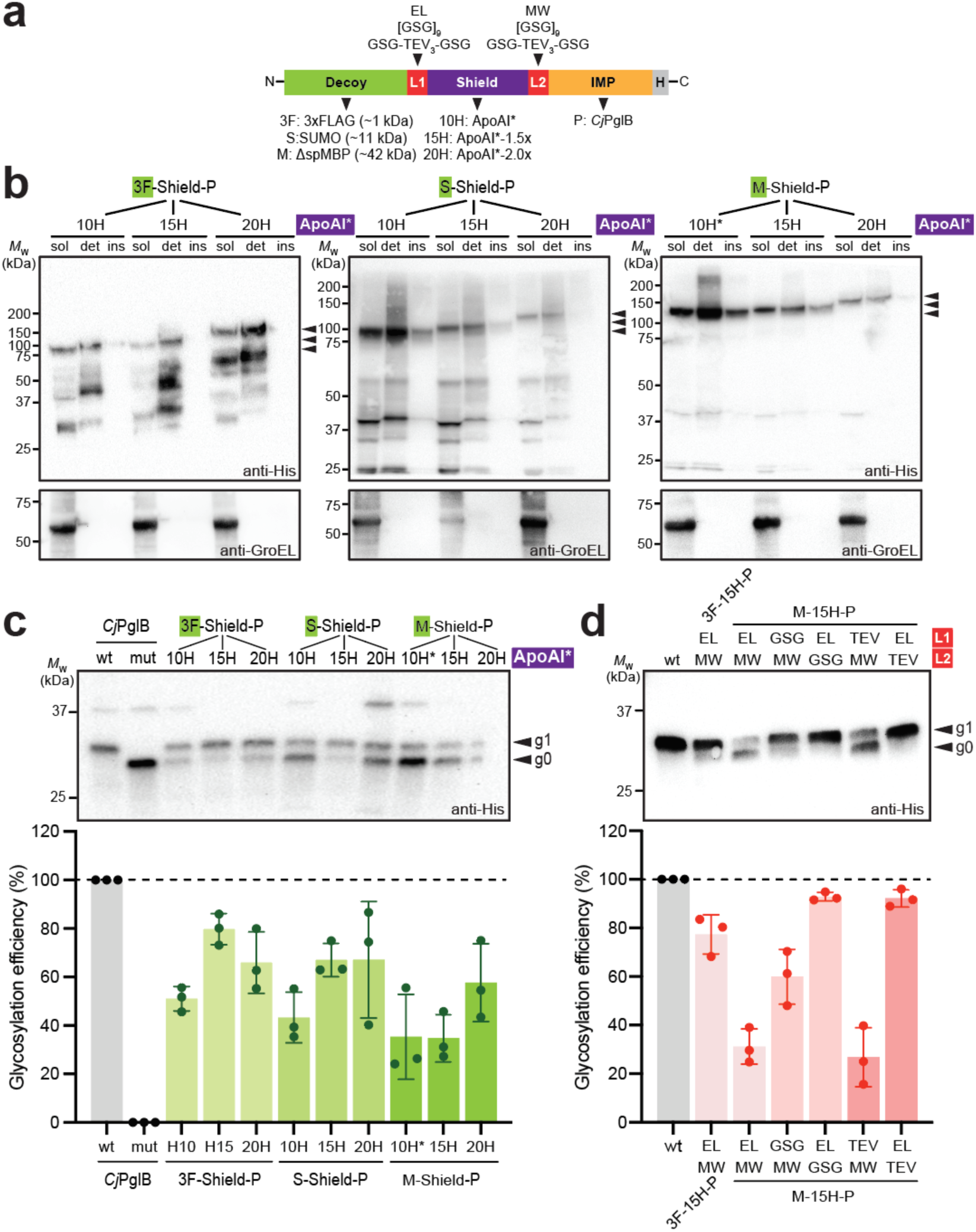
Discovery of highly soluble and highly active SIMPLEx-reformatted *Cj*PglB variants. (a) Schematic of SIMPLEx modularity depicting interchangeable decoy, shield, and linker domains tested in this work. EL: glutamic acid-leucine; MW: methionine-tryptophan; GSG: glycine-serine-glycine linker; TEV: ENLYFQG protease recognition sequence. (b) Immunoblot analysis of soluble (sol), detergent solubilized (det), and insoluble (ins) fractions prepared from BL21(DE3) cells expressing each of the indicated SIMPLEx constructs. An equivalent amount of total protein was loaded in each lane. Blots were probed with anti-His to detect each of the expressed constructs and anti-GroEL antibody to confirm quality of fractionations and equivalent loading of soluble fractions. Black arrows denote full-length expression products. (c) Immunoblot analysis of IVG reactions (top) in which purified scFv13-R4^DQNAT^ acceptor protein was incubated with organic solvent-extracted *Cj*LLOs and SIMPLEx constructs in (b) that were purified from soluble (sol) or detergent solubilized (det) fractions of BL21(DE3) cells. Reactions involving detergent-solubilized wt *Cj*PglB enzyme lacking fusion partners served as positive control (lane 1) while reactions involving detergent-solubilized, catalytically inactive mutant (mut) *Cj*PglB enzyme served as negative control (lane 2). Blots were probed with anti-His to detect the acceptor protein. Black arrows denote aglycosylated (g0) and singly glycosylated (g1) forms of scFv13-R4^DQNAT^. For all immunoblots in (b) and (c), molecular weight (*M*_W_) markers are indicated at left and results are representative of biological replicates (*n* = 3). Glycosylation efficiency (bottom) was determined based on densitometric quantification of the fraction of glycosylated acceptor protein expressed as the ratio g1/[g0+g1]. Data are mean of biological replicates (*n* = 3) ± SD. (d) IVG reactions as in (c) but with SIMPLEx constructs based on 3xFLAG or ΔspMBP decoys fused to ApoAI*-1.5x (15H) with different linker sequences as indicated. IVG reaction involving detergent-solubilized wt *Cj*PglB without any fusions served as positive control (lane 1).

Immunoblot analysis revealed that all newly constructed variants promoted soluble expression of *Cj*PglB without the need for detergents, with similar accumulation patterns as the Sx-*Cj*PglB.v2 construct (**Fig. 2b**). Overall, the larger decoys combined with shorter shields resulted in the highest levels of soluble expression as exemplified by the ΔspMBP and SUMO decoys fused with the 10-helix (10H) ApoAI* domain. It is also worth noting that whereas all fusions involving the ΔspMBP decoy were stably expressed with little visible degradation, the FLAG and SUMO constructs exhibited greater amounts of degradation. Importantly, when the soluble fractions were tested for IVG activity, each construct was observed to catalyze glycan installation on the scFv13-R4^DQNAT^ acceptor protein albeit to varying extents (**Fig. 2c**). Unexpectedly, the constructs that exhibited the most significant solubilization, such as ΔspMBP-10H-*Cj*PglB (a.k.a. Sx-*Cj*PglB.v2) and SUMO-10H-*Cj*PglB that consisted of larger decoys fused with the shortest shield, were the least active in terms of glycosylation efficiency (all <60% efficient). On the other hand, designs involving shorter decoys fused to longer shields were among the most active, with 3xFLAG-15H-*Cj*PglB showing ∼80% glycosylation efficiency.

We suspected that the lower glycosylation efficiency seen for the ΔspMBP-based constructs might be due to interference caused by the relatively bulky ΔspMBP domain. To test this notion, we attempted to create greater spacing and flexibility between the domains of the least active ΔspMBP-15H-*Cj*PglB construct. To this end, we replaced the two-residue linkers (EL and MW) at the decoy:shield and shield:*Cj*PglB junctions with alternative 27-residue linkers, specifically a flexible GlySer linker ([GSG]_9_) or a more rigid TEV protease recognition site-based linker (GSG[ENLYFQG]_3_GSG) (**Fig. 2a**). When either linker was placed between the ΔspMBP decoy and 15H shield, there was only modest improvement in glycosylation efficiency for the GlySer linker and none for the TEV-based linker (**Fig. 2d**). However, when the same linkers were positioned between the 15H shield and *Cj*PglB enzyme, both constructs exhibited a dramatic increase in glycosylation efficiency that reached near 100% and rivaled that of the detergent-solubilized wt *Cj*PglB without any fusion partners. This high glycosylation activity was achieved without compromising the robust soluble expression that was characteristic of the much less active Sx-*Cj*PglB.v2 construct. In fact, the optimized ΔspMBP-EL-15H-[GSG]_9_-*Cj*PglB construct (hereafter Sx-*Cj*PglB.v3) reached 5.6 mg/L when purified directly from the soluble fraction, which was similar to the yield of Sx-*Cj*PglB.v2 purified from the same fraction and >10-fold greater than the yield of wt *Cj*PglB purified from the detergent-solubilized membrane fraction (**Supplementary Fig. 2**). Using the Sx-*Cj*PglB.v3 that was purified from the soluble fraction in IVG reactions resulted in highly efficient glycosylation of the scFv13-R4^DQNAT^ acceptor protein without the need for added detergent, whereas wt *Cj*PglB derived from this same fraction showed no measurable activity (**Supplementary Fig. 3**). Taken together, these results reveal how the straightforward interchangeability of the decoy and shield domains can be used to fine-tune the solubility and activity of SIMPLEx-reformatted IMPs, leading to the identification of catalytically robust designs as exemplified by Sx-*Cj*PglB.v3.

### Machine learning-based design of water-soluble *Cj*PglB variants

In contrast to the universal solubilization afforded by the SIMPLEx methodology, *de novo* designed protein WRAPs (water-soluble RFdiffused amphipathic proteins) are an alternative deep learning-based design approach for custom solubilization of IMP targets ^25^. Similar to ApoAI, WRAPs have a polar exterior and a nonpolar interior complementary to the lipid-facing hydrophobic surface of the target. However, in a notable departure from the SIMPLEx method, each WRAP is computationally tailored to surround the lipid-interacting hydrophobic surfaces of its IMP target using RFdiffusion ^31^ and ProteinMPPN ^32^, generative models for *de novo* protein backbone and sequence design, respectively. Here, we used a similar approach to generate 20 alpha-helical, *Cj*PglB-specific WRAP domains that were designed to render the OST stable and water-soluble without the need for detergents (**Supplementary Fig. 4**). The four highest scoring designs based on pLDDT binder scores (**Supplementary Table 1**) were each genetically fused to the N-terminus of *Cj*PglB with a flexible GlySer linker ([GGGS]_3_) (**Fig. 3a**) and evaluated for soluble expression in the cytoplasm of *E. coli* BL21(DE3) cells. Following purification of the helical WRAP-*Cj*PglB fusion proteins from the soluble fraction of whole-cell lysates, we determined that two of the designs, WRAP-*Cj*PglB.d1 and WRAP-*Cj*PglB.d3, were soluble (**Supplementary Fig. 5**).

**Figure 3.**
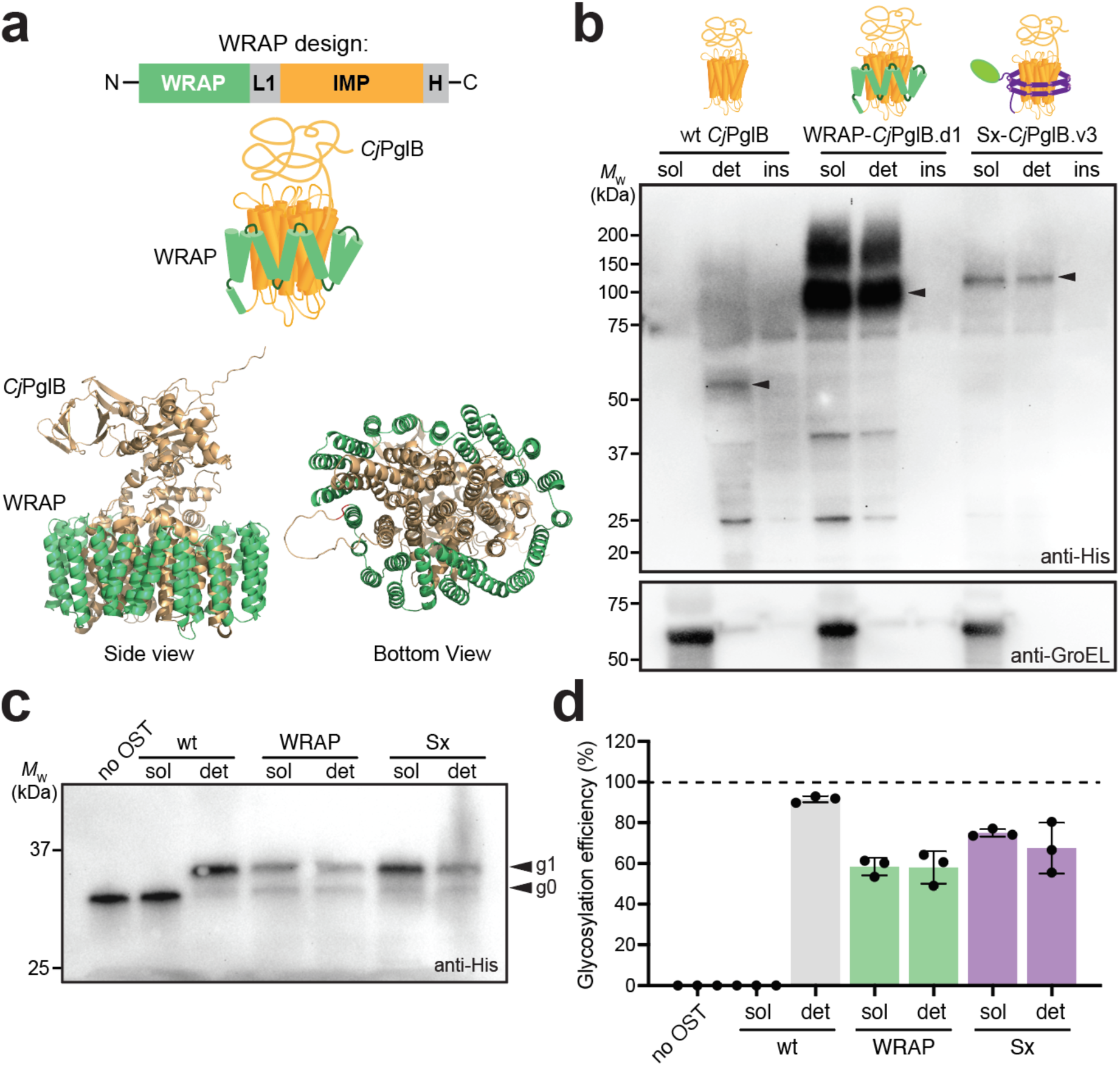
Solubilization of *Cj*PglB using computationally designed helical WRAP domain. (a) Schematic representation (top) and design model (bottom) of WRAP-*Cj*PglB.d1 construct with helical WRAP solubilizing domain (green) custom designed to be complementary to the lipid-facing hydrophobic surface of *Cj*PglB (orange). L1: flexible (GGGS)_3_ linker; H: 10xHis tag. (b) Immunoblot analysis of soluble (sol), detergent solubilized (det), and insoluble fractions (ins) prepared from BL21(DE3) strains expressing wt *Cj*PglB, WRAP-*Cj*PglB.d1, and Sx-*Cj*PglB.v3. Blot was probed with anti-His to detect each of the expressed constructs. An equivalent amount of total protein was loaded in each lane for wt *Cj*PglB and Sx-*Cj*PglB.v3; three times less total protein was loaded for WRAP-*Cj*PglB.d1 samples due to its much higher expression level relative to the other constructs. Black arrows denote full-length expression products. (c) Immunoblot analysis of IVG reactions in which purified scFv13-R4^DQNAT^ acceptor protein was incubated with organic solvent-extracted *Cj*LLOs and *Cj*PglB constructs in (b) that were purified from soluble (sol) or detergent solubilized (det) fractions of BL21(DE3) cells. Reactions involving detergent-solubilized wt *Cj*PglB enzyme lacking fusion partners served as positive control (lane 1) while reactions involving detergent-solubilized, catalytically inactive mutant (mut) *Cj*PglB enzyme served as negative control (lane 2). Blots were probed with anti-His to detect the acceptor protein. Black arrows denote aglycosylated (g0) and singly glycosylated (g1) forms of scFv13-R4^DQNAT^. Molecular weight (*M*_W_) marker is indicated at left and results are representative of biological replicates (*n* = 3). (d) Glycosylation efficiency was determined based on densitometric quantification of the fraction of glycosylated acceptor protein expressed as the ratio g1/[g0+g1]. Data are mean of biological replicates (*n* = 3) ± SD.

Because WRAP-*Cj*PglB.d3 was only weakly detected in the soluble fraction, we turned our attention to WRAP-*Cj*PglB.d1, which showed substantially greater soluble expression. To benchmark the solubility of the WRAP construct, we performed immunoblot analysis of fractionated lysates derived from *E. coli* BL21(DE3) cells expressing either WRAP-*Cj*PglB.d1, unfused wt *Cj*PglB or Sx-*Cj*PglB.v3. This analysis revealed that WRAP-*Cj*PglB.d1 accumulated in the soluble fraction to an extent that greatly exceeded that of Sx-*Cj*PglB.v3 (**Fig. 3b**), demonstrating that the WRAP strategy was highly effective at rendering the 13-TMD OST water-soluble in the cytoplasm without the need for detergents. As before, unfused wt *Cj*PglB partitioned exclusively in the detergent-solubilized membrane fraction. In terms of yield, we recovered 3.3 mg/L of WRAP-*Cj*PglB.d1, which was 6-fold greater than the yield of unfused wt *Cj*PglB but less than the yields for the SIMPLExed *Cj*PglB constructs (**Supplementary Fig. 2**). This latter difference was somewhat surprising given the much stronger expression of the WRAPed construct but could relate to differences in how the enzymes were purified.

To investigate whether WRAPed *Cj*PglB retained the enzymatic function of the native OST, we again turned to the IVG assay using scFv13-R4^DQNAT^ as acceptor protein substrate. Importantly, clear evidence of glycosylation was observed for WRAP-*Cj*PglB.d1 that was purified from the soluble and detergent-solubilized membrane fractions (**Fig. 3c**). This activity was on par with that measured for Sx-*Cj*PglB.v3, with both water-soluble enzymes exhibiting activity that was 60-80% of the activity measured for detergent-solubilized wt *Cj*PglB (**Fig. 3d**). These data establish that our deep learning-based design approach can generate custom alpha-helical WRAP designs that complement the more universal SIMPLEx designs in their ability to create soluble, functionally active versions of *Cj*PglB.

### Kinetic analysis of water-soluble *Cj*PglB variants

Having established the water solubility and glycosylation function of the SIMPLExed and WRAPed versions of *Cj*PglB, we next sought to more rigorously quantify their catalytic performance. To this end, we utilized a peptide substrate-based IVG assay that has been used previously by us and others to establish reaction kinetics for *Cj*PglB and a few of its homologs ^33–36^. This assay is performed with the same solvent-extracted *Cj*LLOs as glycan substrate donor together with a short acceptor peptide substrate containing the DQNAT glycosylation sequon that is terminally labeled with a tetramethylrhodamine fluorophore (5-TAMRA-GS<u>DQNAT</u>F-NH_2_, where the underlined region represents the sequon) (**Fig. 4a**). The extent of TAMRA peptide glycosylation in these reactions is tracked using in-gel fluorescence analysis whereby the increase in molecular weight corresponding to the addition of the ∼1-kDa heptasaccharide is visualized directly in tricine-SDS-PAGE gels. Incubation of these substrates with unfused wt *Cj*PglB, Sx-*Cj*PglB.v3, or WRAP-*Cj*PglB.d1 resulted in the formation of fluorophore-labeled glycopeptide products dependent on the fraction from which the OST was purified (**Supplementary Fig. 6a**). Specifically, whereas all three enzymes glycosylated the TAMRA-peptide when the enzymes were purified from the detergent-solubilized membrane fraction, only the Sx-*Cj*PglB.v3 and WRAP-*Cj*PglB.d1 enzymes were able to catalyze TAMRA peptide glycosylation when the enzymes were purified from the soluble fraction, consistent with our IVG results using scFv13-R4^DQNAT^ as acceptor substrate. These water-soluble OSTs were determined to be 85% and 30% efficient, respectively, while unfused wt *Cj*PglB purified from the detergent-solubilized membrane fraction was 91% efficient (**Supplementary Fig. 6b**).

**Figure 4.**
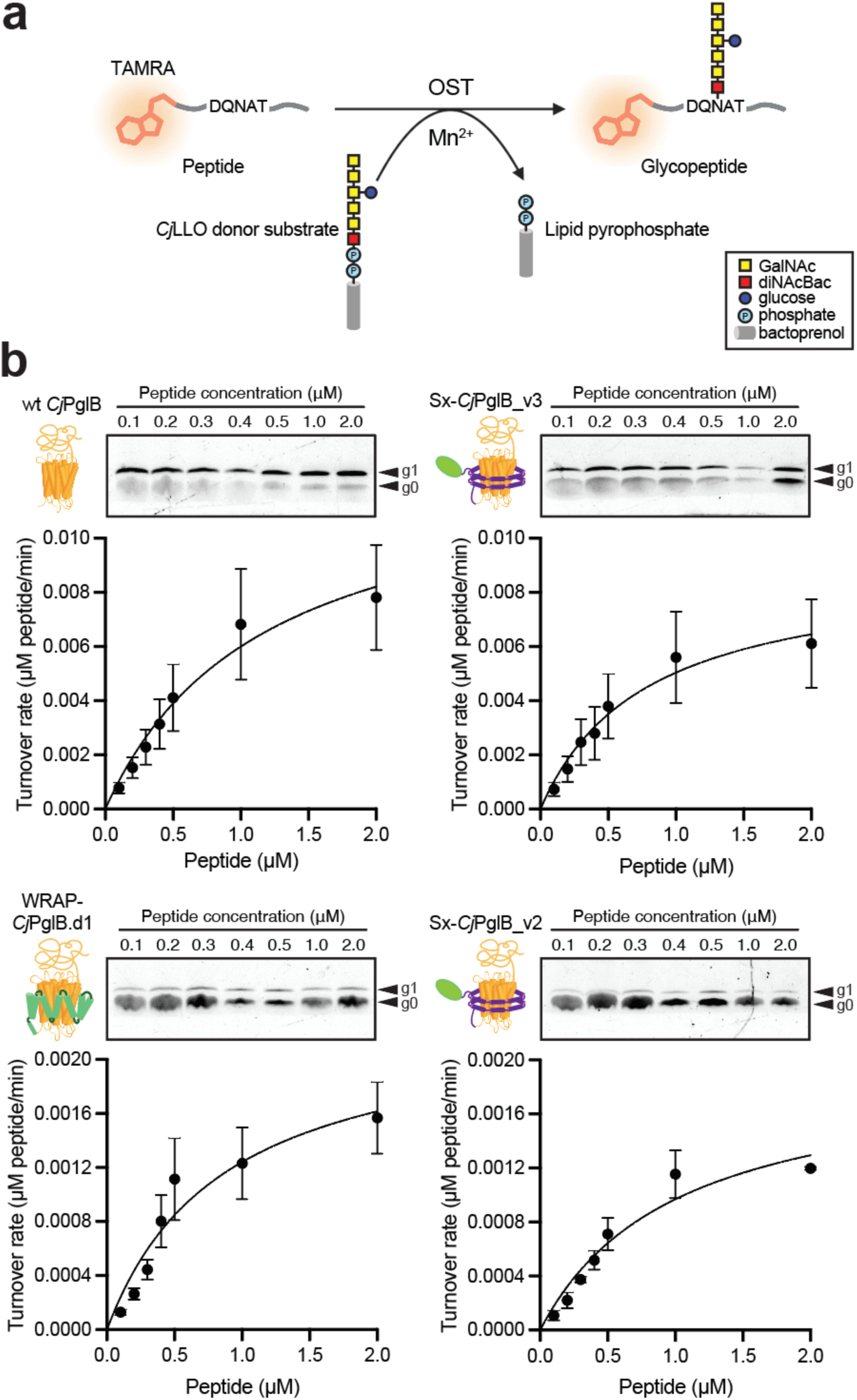
Glycosylation reaction kinetics of water-soluble OSTs. (a) Schematic of TAMRA peptide IVG assay. Incubation of TAMRA-labeled peptides with OSTs, *Cj*LLOs, and divalent metal ions (e.g., Mn^2+^) results in transfer of heptasaccharide glycans to form glycopeptide products. (b) Tricine SDS-PAGE analysis of IVG reactions with different amounts of TAMRA-labeled peptide and 0.5 μM of each OST. Graph at right depicts determination of Michaelis–Menten kinetics using IVG reaction data. Data fitting was by nonlinear regression analysis according to Michaelis–Menten model using Prism 10 for MacOS (version 10.3.0). Black arrows denote aglycosylated (g0) and singly glycosylated (g1) forms of TAMRA-labeled peptides. Tricine SDS-PAGE gels in each panel are representative of three biological replicates. Data in corresponding graphs are average of biological replicates (*n* = 3) ± SD.

Using this TAMRA peptide glycosylation assay, we proceeded to estimate the kinetic constants for all three OSTs. Specifically, Michaelis-Menten kinetics were determined by performing IVG reactions with increasing concentrations of the TAMRA peptide substrate and quantifying the in-gel fluorescence intensities of the aglycosylated (g0) and glycosylated (g1) bands as a function of peptide concentration (**Fig. 4b**) and used these data to determine *K_m_* and *V*_max_ values (**Table 1**). From this analysis, the water-soluble Sx-*Cj*PglB.v3 and detergent-solubilized wt *Cj*PglB constructs exhibited comparable kinetics, clearly outperforming the WRAP-*Cj*PglB.d1 construct (**Table 1**). One possible explanation for this discrepancy is that WRAPs are designed to bind tightly to their IMP target, which in turn could affect their catalytic activity. However, we also note that whereas Sx-*Cj*PglB.v3 underwent significant optimization of its linker and decoy domains to improve its activity, the WRAP-*Cj*PglB.d1 construct was not subjected to any such optimization. Thus, a fairer comparison is between the straight out-of-the-box WRAP-*Cj*PglB.d1 construct and its non-optimized Sx-*Cj*PglB.v2 counterpart. Indeed, the kinetics of these two constructs was similar, with WRAP-*Cj*PglB.d1 slightly outperforming Sx-*Cj*PglB.v2 (**Figure 4b** and **Table 1**).

**Table 1.** Kinetic parameters for wild-type and water-soluble *Cj*PglB enzymes.

| OST | $K_M$ ($\mu\text{M}$) | $V_{\max}$ (peptide $\text{h}^{-1}$) | Turnover rate ($\text{h}^{-1}$) |
| --- | --- | --- | --- |
| wt CjPglB | $1.2 \pm 0.18$ | $0.78 \pm 0.26$ | $0.072 \pm 0.001$ |
| Sx-CjPglB.v2 | $1.0 \pm 0.24$ | $0.12 \pm 0.01$ | $0.023 \pm 0.002$ |
| Sx-CjPglB.v3 | $0.78 \pm 0.14$ | $0.54 \pm 0.16$ | $0.056 \pm 0.003$ |
| WRAP-CjPglB.d1 | $0.87 \pm 0.20$ | $0.14 \pm 0.03$ | $0.044 \pm 0.006$ |

We next decided to determine and compare initial turnover rates because impractically high peptide concentrations would be required to reach *V*_max_, as was noted previously ^33^. Specifically, the turnover rate for all four OSTs was determined from the initial, linear range of time course data (**Supplementary Fig. 7**). This analysis revealed a similar trend in terms of catalytic performance with the turnover rates decreasing in the order: wt *Cj*PglB > Sx-*Cj*PglB.v3 > WRAP-*Cj*PglB.d1 > Sx-*Cj*PglB.v2. Collectively, these kinetic data help to explain the differences in glycosylation efficiency achieved by these enzymes in the context of the scFv13-R4^DQNAT^ acceptor substrate while also providing further confirmation that our water-soluble *Cj*PglB designs retained the catalytic function of the native OST without the need for detergents. It is also worth noting that the kinetic constants determined here compared favorably with values reported previously for detergent-solubilized wt *Cj*PglB assayed with LLOs derived from either chemical synthesis or cellular biosynthesis ^34, 35^.

## DISCUSSION

In this work, we demonstrate that even the most topologically complex GTs—those belonging to the GT-C structural superfamily—can be systematically converted into functional, water-soluble enzymes using rational and computational protein engineering approaches. By applying both SIMPLEx, a universal solubilization platform, and WRAPs, a custom deep learning–guided design strategy, we overcame the longstanding challenge of producing multi-pass membrane glycosyltransferases in a tractable, detergent-free format. The successful reformatting of *Cj*PglB, an archetypal GT-C enzyme containing 13 transmembrane helices, highlights the robustness of these platforms and provides a blueprint for solubilizing other GT-C family members whose structures and biochemical properties remain poorly understood.

Importantly, both strategies produced water-soluble PglB variants that retained the defining catalytic hallmarks of wild-type enzyme function. SIMPLEx-solubilized *Cj*PglB performed with kinetic parameters and glycosylation efficiencies comparable to detergent-solubilized wild type, and WRAP-designed variants achieved similar functional outcomes despite being built from entirely distinct protein design principles. The convergence of function across such divergent engineering approaches underscores a key insight: many of the topological requirements for membrane integration can be decoupled from the catalytic machinery itself, provided that the hydrophobic surfaces of the enzyme are adequately shielded and its dynamic architecture is preserved.

Together, these results establish SIMPLEx and WRAPs as versatile and complementary strategies for overcoming the expression and solubility bottlenecks that have constrained molecular characterization of the GT-C superfamily. While SIMPLex is universally applicable, WRAPs require custom protein design, which in turn contributes to stability and enables structure determination. As structural data for GT-C enzymes remain scarce, the ability to generate high-yield, monodisperse, detergent-free variants creates new opportunities for biochemical, biophysical, and structural interrogation, including crystallography, cryo-EM, and deep mutational scanning. Moreover, these approaches could be extended beyond GT-Cs to other polytopic integral membrane enzymes whose inherent hydrophobicity currently limits mechanistic study and biotechnological deployment.

Despite these advances, several open questions remain. Future efforts will be needed to determine how solubilization affects long-range conformational dynamics of GT-C enzymes, particularly those linked to membrane-dependent substrate binding or conformational gating. Additionally, while both design approaches preserved catalytic activity toward model substrates, broader analyses will be required to assess how well substrate specificity, glycan preferences, or in vivo selectivity are maintained across diverse donor and acceptor contexts. Finally, the full generalizability of these solubilization strategies across the phylogenetic diversity of GT-C enzymes—including eukaryotic STT3 paralogs—remains an exciting direction for future exploration.

Overall, our work provides foundational evidence that structurally complex membrane-integrated glycosyltransferases can be reformatted into stable, active, water-soluble catalysts. By enabling high-yield expression, functional characterization, and integration into cell-free and synthetic glycosylation platforms, SIMPLEx and WRAPs open new avenues for discovery and application across glycobiology, enzymology, and glycoengineering.

## MATERIALS AND METHODS

### Bacterial strains and plasmids

*E. coli* strain NEB5α (New England Biolabs), a derivative of DH5α, was used for all plasmid construction, maintenance, and DNA purification. *E. coli* strain BL21(DE3) (Novagen) was used for all protein expression experiments. Plasmid pSN18 ^37^ was used for expression of wt *Cj*PglB, while plasmid pET28a-scFv13-R4(N34L, N77L)^DQNAT-His^^6^ ^36^ was used for expression of the scFv13-R4^DQNAT^ acceptor protein. *E. coli* strain CLM24 carrying plasmid pMW07-pglΔB ^38^, which encodes the entire *C. jejuni pgl* pathway except for *Cj*PglB ^39^, was used for preparing *Cj*LLOs. CLM24 is a W3110 derivative with a deletion in the O-antigen ligase WaaL that prevents glycan transfer to lipid A-core ^40^, enabling accumulation of the *C. jejuni* heptasaccharide *N-*glycan on undecaprenyl pyrophosphate (Und-PP).

To express *Cj*PglB in the original SIMPLEx format, plasmid pET28a-Sx-*Cj*PglB.v1 was constructed by PCR amplifying the gene encoding *Cj*PglB from pSN18 and inserting it between ΔspMBP and ApoAI* in plasmid pET28(+)-ΔspMBP-(*Nde*I)-GT-(*Eco*RI)-ApoAI* ^16^ using Gibson Assembly ^41^. To express *Cj*PglB in the redesigned SIMPLEx format, plasmid pBAD-Sx-*Cj*PglB.v2 encoding *E. coli* MBP, human ApoAI*, and *Cj*PglB in the pSN18 backbone was constructed commercially (Integrated DNA Technologies). The resulting plasmid placed the Sx-*Cj*PglB.v2 construct under control of an arabinose-inducible promoter and introduced unique restriction sites flanking each of the individual genes and a C-terminal 10xHis tag as follows: pBAD-(*Spe*l)-ΔspMBP-(*Sac*I)-ApoAI*-(*Nco*I)-*Cj*PglB-(*Eco*RI)-10xHis. The plasmids pBAD-ΔspMBP-*Cj*PglB and pBAD-ApoAI*-*Cj*PglB for expressing control constructs lacking either the ApoAI* or ΔspMBP domains, respectively, were constructed by PCR amplification of the complete pBAD-Sx-*Cj*PglB.v2 plasmid excluding either the ApoAI* or ΔspMBP sequence, followed by self-ligation of the linear PCR product in *E. coli* NEB5α cells. To generate SIMPLEx variants with alternative decoy domains, DNA encoding 3xFLAG and SUMO domains was synthesized (Integrated DNA Technologies), amplified by PCR, and subsequently used to replace the DNA encoding the ΔspMBP domain in pBAD-Sx-*Cj*PglB.v2 by Gibson Assembly. Likewise, to introduce longer ApoAI* domains consisting of 15 and 20 helices, DNA encoding the 15H and 20H helices was synthesized (Integrated DNA Technologies), PCR amplified, and used to replace the 10-helix (10H) ApoAI* domain by Gibson Assembly. To introduce alternative linkers, DNA encoding the flexible GlySer linker ([GSG]_9_) and the TEV protease recognition site-based linker (GSG[ENLYFQG_3_GSG) were synthesized (Integrated DNA Technologies), PCR amplified and ligated into plasmid pBAD-Sx-*Cj*PglB.v2 at either the *Sac*I or *Nco*I restriction site. To express the WRAP-*Cj*PglB designs, DNA encoding the top four WRAP designs was codon optimized for expression in *E. coli* and synthesized (Twist Bioscience). The resulting synthetic DNA was PCR amplified and inserted upstream of the gene encoding wt *Cj*PglB in pSN18 by Gibson Assembly, resulting in plasmids pBAD-WRAP-*Cj*PglB.d1 to v4. All newly constructed plasmids were generated using standard molecular cloning techniques and verified by Oxford nanopore whole plasmid sequencing (Eurofins Genomics) or Sanger sequencing at the Genomics Facility of the Cornell Biotechnology Resource Center.

### Design of protein WRAPs

Initial WRAPs were generated parametrically using sushimaki (https://github.com/davidekim/sushimaki), a python script that uses DeepTMHMM ^42^ to predict the membrane spanning residues and topology of the target used to estimate the radius (*tr*), height (*th*), and central axis of the transmembrane region, and PyRosetta ^43^ to generate and place the helical WRAP around the transmembrane region of the target. The main purpose of the script is to roughly generate WRAPs around a given target to use as inputs for structure refinement using partial RFdiffusion ^31, 44^. Ideal helices of *hn* residues per helix are generated, where *hn* is determined by *th,* and the number of helices (*n*) is determined by *tr* using the following equations, respectively:

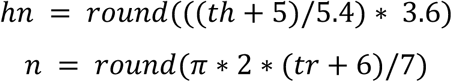

*hn* is calculated based on the rise per residue of an ideal helix, 5.4/3.6, plus a 5-angstrom buffer added to *th*. *n* is determined from the circumference of the WRAP divided by 7, a value that was empirically derived from WRAPs in a previous study ^25^. *hn* and *n* can also be provided as command line options where *th* and *tr* are calculated using the same equations rather than estimated by DeepTMHMM. To generate the WRAP, sushimaki places a helix approximately along the transmembrane central axis by superimposing the center of masses of the top and bottom 4 residue CA atoms and all CAs of the helix to the center of masses of the top and bottom 2 residue CAs of each membrane spanning chain segment and all spanning CAs of the transmembrane region, respectively. The aligned helix is then translated perpendicular to the transmembrane central axis at a distance of *tr* plus 6 angstroms to roughly accommodate side chain packing. The helix up and down orientation is flipped if the N or C terminus is on the opposite side of the C or N terminus of the target, respectively. The helix is then rotated around the central axis in 5 degree increments until the distance between the respective termini is at the minimum to enable fusion of the WRAP to the target with a short flexible linker. The remaining helices are generated as copies of the helix and then placed evenly around the transmembrane region by rotating around the central axis with alternating up and down orientations. The helices are then connected by 3 residue loops into a single chain using the PyRosetta loop modeler. Both forward and reverse rotating WRAPs with –30, 0, and 30 degree rotation offsets are generated for N and C terminal target fusions for a total of 12 WRAPs per *n* and *hn* parameter values used. For the target protein, 5OGL, WRAPs were generated using *n* values of 19 to 24 helices and *hn* values of 16 and 17 residues to sample a larger range of WRAP structures verses *n* = 19 and *hn* = 17 derived from DeepTMHMM for 5OGL.

The WRAPed target complexes were then refined using partial RFdiffusion conditioned on the target structure with “diffuser.partial_T” of 30 and “diffuser.T” of 50 steps total to generate 1440 designs. These partially diffused backbones were then sequence designed and validated by a protein-protein interaction design pipeline described previously ^45^ involving SolubleMPNN ^32^, Rosetta FastRelax ^46^, and Alphafold2 (AF2) ^47^ initial guess. Briefly described here, the pipeline uses SolubleMPNN to design sequences for the partially diffused WRAP backbones while keeping the target sequence fixed. These designs are also Rosetta FastRelaxed and sequence re-designed. The structures of the designed complexes are then predicted using AF2 initial guess. 8 designs were selected with binder (WRAP) pLDDT > 80, interface PAE (PAE_i) < 10, and SAP (spatial aggregation propensity) ^48^ < 600 from 2880 AF2 models. To further improve these metrics, an iterative partial diffusion optimization strategy (https://github.com/davidekim/ppi_iterative_opt) was used for each of the 8 selected designs involving iterations of partial diffusion, SolubleMPNN plus Rosetta FastRelax, and AF2 validation, where the design with the lowest PAE_i in each iteration is used for the next iteration. Up to 10 iterations were run, and 5 final designs with the highest pLDDTs were selected from the top 20 designs with the lowest PAE_i values from 13,053 designs. 4 designs with exposed hydrophobic transmembrane target residues were excluded upon visual inspection.

### Subcellular fractionation

For recombinant expression of wt *Cj*PglB and the water-soluble *Cj*PglB variants, *E. coli* BL21(DE3) cells harboring the appropriate expression plasmid were cultured overnight at 37 °C in 10 mL Luria Bertani (LB) medium supplemented with 100 μg/mL ampicillin. Overnight cells were subcultured into 1 L of Terrific Broth (TB) containing the same antibiotic and grown at 37 °C with shaking until the optical density at 600 nm (OD_600_) reached ≈ 0.7. Protein expression was induced by addition of L-arabinose to a final concentration of 0.2% (v/v), and cultures were incubated at 16 °C overnight (∼16–20 h). Cells were harvested by centrifugation at 5,000 ×g for 15 min at 4 °C and resuspended in Buffer A (50 mM HEPES, 250 mM NaCl, pH 7.4; sterile-filtered). Cell lysis was performed using an EmulsiFlex-C5 high-pressure homogenizer (Avestin). The lysates were cleared by centrifugation at 3,000 ×g for 20 min at 4 °C to remove cellular debris. The clarified lysates were subjected to ultracentrifugation at 100,000 ×g for 2 h at 4 °C to fractionate soluble and membrane-associated proteins. The supernatant from the ultracentrifugation step was collected as the soluble fraction. The resulting pellet was resuspended in Buffer B (50 mM HEPES, 250 mM NaCl, 1% (w/v) n-dodecyl-β-D-maltoside (DDM), pH 7.4; sterile-filtered) and incubated at room temperature for 1 h to extract membrane proteins. Following detergent solubilization, samples were again centrifuged at 100,000 ×g for 1 h at 4 °C. The resulting supernatant was collected as the detergent-solubilized membrane fraction, while the remaining pellet was resuspended in Buffer B and defined as the insoluble fraction. Whole-cell lysate samples (i.e., lysates not subjected to ultracentrifugation) were retained as whole cell fractions for comparative analysis. **Protein purification.** *Cj*PglB proteins were purified from the soluble fraction of the lysate, the detergent-solubilized fraction of the lysate, or the whole-cell fraction using immobilized metal affinity chromatography (IMAC). For purification from whole-cell and soluble fractions, all buffers were prepared using Buffer A, while purification from detergent-solubilized fractions involved buffers prepared using Buffer B. Ni-NTA agarose resin (ThermoFisher) was equilibrated in the appropriate buffer by washing with five column volumes (CVs). Resin was added to the prepared fractions and incubated at 4 °C with gentle agitation. For wild-type *Cj*PglB purified from the detergent-solubilized fraction, incubation was performed overnight at 4 °C. All other incubations were performed for 1 h at 4 °C. After binding, the resin was washed with 20 mM imidazole-containing wash buffer prepared from either Buffer A or B. Bound enzymes were eluted with elution buffer supplemented with 300 mM imidazole in the same base buffer. Elution was monitored by absorbance at 280 nm using a spectrophotometer (NanoDrop) at 1-mL intervals, and fractions were pooled until the absorbance signal approached baseline. Eluted proteins were buffer exchanged into either Buffer A or B using centrifugal concentrators (ThermoFisher) prior to downstream analysis.

For final yield determination and kinetic studies, all *Cj*PglB proteins were further purified via either affinity chromatography using amylose resin for SIMPLEx constructs or size-exclusion chromatography (SEC) for wt and WRAPed *Cj*PglB constructs. Amylose purification was performed by 10-fold dilution of IMAC-eluted samples into equilibration buffer (20 mM Tris-HCl, 200 mM NaCl, 1 mM EDTA, pH 7.2). Pre-washed amylose resin (New England Biolabs) was added and incubated with samples for 1 h at 4 °C. After washing, bound proteins were eluted with equilibration buffer supplemented with 20 mM maltose, and dialyzed into Buffer A overnight. SEC was performed using a Superdex 200 Increase column (Cytiva) pre-equilibrated in Buffer A (for WRAP-*Cj*PglB.d1) or Buffer B (for wt *Cj*PglB). Eluted fractions were collected and analyzed based on chromatographic profiles. Purity assessments were conducted by SDS-PAGE analysis followed by Coomassie staining of gels.

The scFv13-R4^DQNAT^ acceptor protein was purified from whole-cell lysates by IMAC. Briefly, BL21(DE3) cells carrying pET28a-scFv13-R4(N34L, N77L)^DQNAT-His6^ were initially grown overnight in 10 mL of LB supplemented with 50 μg/mL kanamycin, subcultured into 1 L of LB with the same antibiotic on the following day, and induced with 0.1 mM IPTG at 20 °C overnight. Cells were harvested and resuspended in Buffer C (20 mM Tris-HCl, 250 mM NaCl, pH 7.2; sterile-filtered) and lysed using an EmulsiFlex-C5 high-pressure homogenizer (Avestin). Lysates were clarified by centrifugation at 3,000 ×g for 20 min at 4 °C to remove cell debris. Ni-NTA agarose resin (ThermoFisher) was equilibrated in Buffer C and incubated with clarified lysates for 1 h at 4 °C with gentle agitation. After binding, the resin was washed with Buffer C supplemented with 20 mM imidazole with five CVs. Bound proteins were eluted with Buffer C supplemented with 300 mM imidazole. Elution was monitored at 280 nm using a spectrophotometer (NanoDrop) in 1-mL fractions, which were pooled until the absorbance signal approached baseline. Eluted proteins were buffer exchanged into Buffer C using PD-10 desalting columns (Cytiva) according to the manufacturer’s instructions.

### Coomassie staining and immunoblotting

Protein samples were denatured by addition of sodium dodecyl sulfate–polyacrylamide gel electrophoresis (SDS–PAGE) loading buffer containing 10% β-mercaptoethanol, followed by heating at 90 °C for 10 min. Electrophoretic separation was performed using either precast 4–20% Tris-Glycine gels (Invitrogen) or AnyKD Mini-PROTEAN TGX gels (Bio-Rad). Gels from Bio-Rad were specifically used for scFv13-R4^DQNAT^ samples, while Invitrogen gels were used for all other protein samples. For total protein visualization, gels were stained using Coomassie Brilliant Blue G-250 (Bio-Rad) following the manufacturer’s instructions. Stained gels were imaged using a standard flatbed scanner or gel documentation system. For immunoblotting, following electrophoresis, proteins were transferred from SDS-PAGE gels onto polyvinylidene difluoride (PVDF) membranes (Immobilon-P, Millipore) using a wet tank transfer system (ThermoFisher), following the manufacturer’s protocol. Membranes were blocked for 1 h at room temperature or overnight at 4 °C in TBST buffer (20 mM Tris, 150 mM NaCl, 0.05% Tween-20) containing 5% (w/v) non-fat dry milk. Membranes were then washed five times with TBST for 5 min before primary antibody incubation. The following antibodies were used for immunoblotting: polyhistidine (6x-His) tag-specific polyclonal antibody (Abcam, cat # ab9108; diluted 1:10,000), *E. coli* GroEL-specific antibody (Sigma-Aldrich, cat # G6532; diluted 1:10,000), and donkey anti-rabbit IgG conjugated to horseradish peroxidase (HRP) (Abcam, cat # ab7083; diluted 1:10,000). After probing with primary and second antibodies, the membranes were washed five times with TBST for 10 min. Chemiluminescence was developed using ECL substrate (Bio-Rad), and signals were imaged using a ChemiDoc MP Imaging System (Bio-Rad). Glycosylation efficiency was determined by performing densitometry analysis of protein bands in anti-His immunoblots using ImageJ software ^49^ as described at https://imagej.net. Briefly, bands corresponding to g0 in each lane were grouped as a row or a horizontal “lane” and quantified using the gel analysis function in ImageJ. The bands corresponding to g1 were analyzed identically. The resulting intensity data for g0 and g1 was used to calculate percent glycosylated expressed according to the following ratio: g1/[g0+g1]. Efficiency data was calculated from immunoblots corresponding to three biological replicates, with all data reported as the mean ± SD. Statistical significance was determined by paired Student’s *t* tests (\**p* < 0.05, \*\**p* < 0.01; \*\*\**p* < 0.001; \*\*\*\**p* < 0.0001) using Prism 10 for MacOS version 10.3.0.

### *In vitro* glycosylation (IVG)

For *in vitro* glycosylation of scFv13-R4^DQNAT^, 50 μL of *in vitro* glycosylation buffer (50 mM HEPES (pH 7.4), 25 mM MnCl_2_, and 0.1% (w/v) DDM) was supplemented with 1 μg of scFv13-R4^DQNAT^ acceptor protein, 1–20 μg of purified detergent-solubilized wt *Cj*PglB or detergent-free water-soluble *Cj*PglB, and 5–10 μL of solvent-extracted *Cj*LLOs (∼10–20% (v/v) of the total reaction volume) and incubated at 30 °C for 16–20 h. Enzyme amounts were normalized based on molar concentrations rather than total protein mass to ensure accurate comparison of catalytic efficiencies. Organic solvent extraction of *Cj*LLOs bearing the *C. jejuni* heptasaccharide glycan from the membrane of *E. coli* cells was performed as follows. *E. coli* CLM24 cells carrying plasmid pMW07-pglΔB were cultured overnight in LB medium supplemented with 20 μg/mL chloramphenicol at 37 °C. Overnight cultures were subcultured into 1 L of TB medium with chloramphenicol and grown at 37 °C until the OD_600_ reached ≈ 0.8. Protein expression was induced with 0.2% (v/v) L-arabinose, and cultures were incubated at 30 °C overnight. The next day, cells were harvested by centrifugation at 5,000 ×g for 15 min at 4 °C, resuspended in methanol, and transferred onto glass plates for drying overnight in a fume hood. The dried cell pellets were mechanically pulverized using a glass rod and subjected to lipid extraction with a 2:1 (v/v) chloroform:methanol mixture. Samples were sonicated in a water bath for 10 min and centrifuged at 3,000 ×g for 10 min at 4 °C. The supernatant was decanted, and the pellet was washed with water, followed by additional sonication and centrifugation. The resulting pellet was then extracted with a chloroform:methanol:water (10:10:3, v/v/v) mixture, sonicated again, and centrifuged at 3,000 ×g for 10 min at 4 °C. The upper aqueous phase was discarded, and the lower organic layer was collected into a fresh conical tube. After a final centrifugation at 3,000 ×g for 10 min at 4 °C, the organic phase was transferred to a clean glass plate and dried overnight in a fume hood. Dried *Cj*LLOs were resuspended in a detergent-based buffer (50 mM HEPES, 0.1% (w/v) DDM, pH 7.4) and stored at −20 °C until further use in glycosylation assays.

### Determination of enzyme kinetics

Enzyme kinetics were determined by performing IVG with TAMRA-labeled peptides as acceptor substrates. For turnover rate experiments, each reaction was prepared in a total volume of 100 µL *in vitro* glycosylation buffer supplemented with 1–2 μM of purified detergent-solubilized wt *Cj*PglB or detergent-free water-soluble *Cj*PglB, 5–10 μL (∼10–20% (v/v)) of solvent-extracted *Cj*LLOs, 0.5 µM of 5-TAMRA-GSDQNATF-NH_2_ peptide (GenScript), and ddH_2_O as needed. Reactions were incubated in a water bath at 30 °C with samples collected at different time points and stopped by boiling at 90 °C for 5 min. For Michaelis–Menten kinetics, reactions were performed in a total volume of 10 µL containing: 1 µL of *in vitro* glycosylation buffer, 0.5 μM of purified detergent-solubilized wt *Cj*PglB or detergent-free water-soluble *Cj*PglB, 5–10 μL (∼10–20% (v/v)) of solvent-extracted *Cj*LLOs, varying concentrations of 5-TAMRA-GSDQNATF-NH_2_ (ranging from 0.1–2.0 μM), and ddH_2_O as needed. The reactions were incubated for 90 min at 30 °C and stopped by boiling the sample at 90 °C for 5 min.

All reaction products were subjected to in-gel fluorescence analysis by diluting samples in Novex Tricine SDS Running Buffer (1x), mixing with dye that was produced in-house, and boiling at 80 °C for 2 min. The dye consisted of 200 mM Tris-Cl (pH 6.8), 8% (w/v) SDS (electrophoresis grade), and 40% (v/v) glycerol. Prior to gel loading, samples were diluted in Novex Tricine SDS Running Buffer (1x) so that concentration of total peptide in each lane was identical. A total of 5 µL of each sample was loaded onto Novex 16% Tricine Mini Protein Gels (1.0 mm thickness). The Spectra™ Multicolor Low Range Protein Ladder was used as the molecular weight marker. The gel was run at 70 V for 1.5 h at 4 °C to separate glycosylated peptides (g1) from non-glycosylated peptides (g0), and bands were subsequently visualized by fluorescence scanning of gels (ex: 488 nm; em: 526 nm) using a ChemiDoc MP Imaging System (Bio-Rad). DyLight 550 was used to visualize the fluorescently labeled peptides, while the Spectra ladder was visualized using Cy5.5. Using Prism 10 for MacOS (version 10.3.0), reaction velocities for each OST were determined by nonlinear regression analysis according to the Michaelis–Menten model while initial turnover rates were determined by linear regression analysis of the linear portion of the time-course reactions.

## Acknowledgements

This work was supported by the Defense Advanced Research Projects Agency (DARPA contract W911NF-23-2-0039 to M.C.J. and M.P.D.), the Defense Threat Reduction Agency (grants HDTRA1-15-10052 and HDTRA1-20-10004 to M.P.D. and M.C.J.), the National Science Foundation (grants CBET-1605242 to M.P.D., CBET-1936823 and MCB-1413563 to M.P.D. and M.C.J.), and the National Institutes of Health (grants R01GM127578 to M.P.D.). L.M. is funded by HHMI Helen Hay Whitney postdoctoral fellowship. K.K. was supported by an American Heart Association Postdoctoral Fellowship. T.C.D. was supported by a Hartwell Foundation Postdoctoral Fellowship. E.J.B was supported by an NIH/NIGMS Chemical Biology Interface Training Grant (T32GM138826) and an NSF Graduate Research Fellowship (DGE-2139899). S.W.H. was supported by a training grant from the National Institutes of Health NIBIB (T32EB023860) and a Cornell Founder’s Dissertation Fellowship. M.C. was supported by the National Science Foundation (DBI-2244288) through the Rosetta Commons REU. TJ was supported by an EMBO postdoctoral fellowship (ALTF 336-2021) and the Danish National Research Foundation (DNRF107).

## Author Contributions

**Y.H.K.:** conceptualization, methodology, validation, formal analysis, investigation, writing – original draft, writing – review & editing, visualization. **L.M.:** conceptualization, methodology, validation, formal analysis, investigation, writing – original draft, writing – review & editing, visualization. **K.K.:** methodology, validation, formal analysis, investigation related to cloning, expression and purification of SIMPLEx constructs. **D.E.K.:** methodology and investigation related to WRAP designs, writing – original draft, writing – review & editing. **T.C.D.:** methodology and investigation related to mass spectrometry analysis. **E.J.B.:** methodology and investigation related to mass spectrometry analysis. **C.K.B.:** methodology and investigation related to mass spectrometry analysis. **B.S.:** methodology and investigation related to enzyme kinetics. **S.W.H.:** methodology and investigation related to plasmid construction. **K.A.M.:** methodology and investigation related to conjugate vaccine expression and glycosylation. **A.T.:** methodology and investigation related to conjugate vaccine expression and glycosylation. **M.C.:** methodology and investigation related to WRAP designs. **D.M.:** conceptualization, methodology and investigation related to SIMPLEx designs. **T.J.:** conceptualization, methodology and investigation related to SIMPLEx designs. **H.C.:** supervision. **M.C.J.:** conceptualization, supervision, funding acquisition. **D.B.:** conceptualization, supervision, funding acquisition. **M.P.D.:** conceptualization, methodology, formal analysis, writing – original draft, writing – review & editing, visualization, supervision, project administration, funding acquisition.

