## Supplementary Information for "Topological reprogramming transforms an integral membrane oligosaccharyltransferase into a water-soluble glycosylation catalyst"

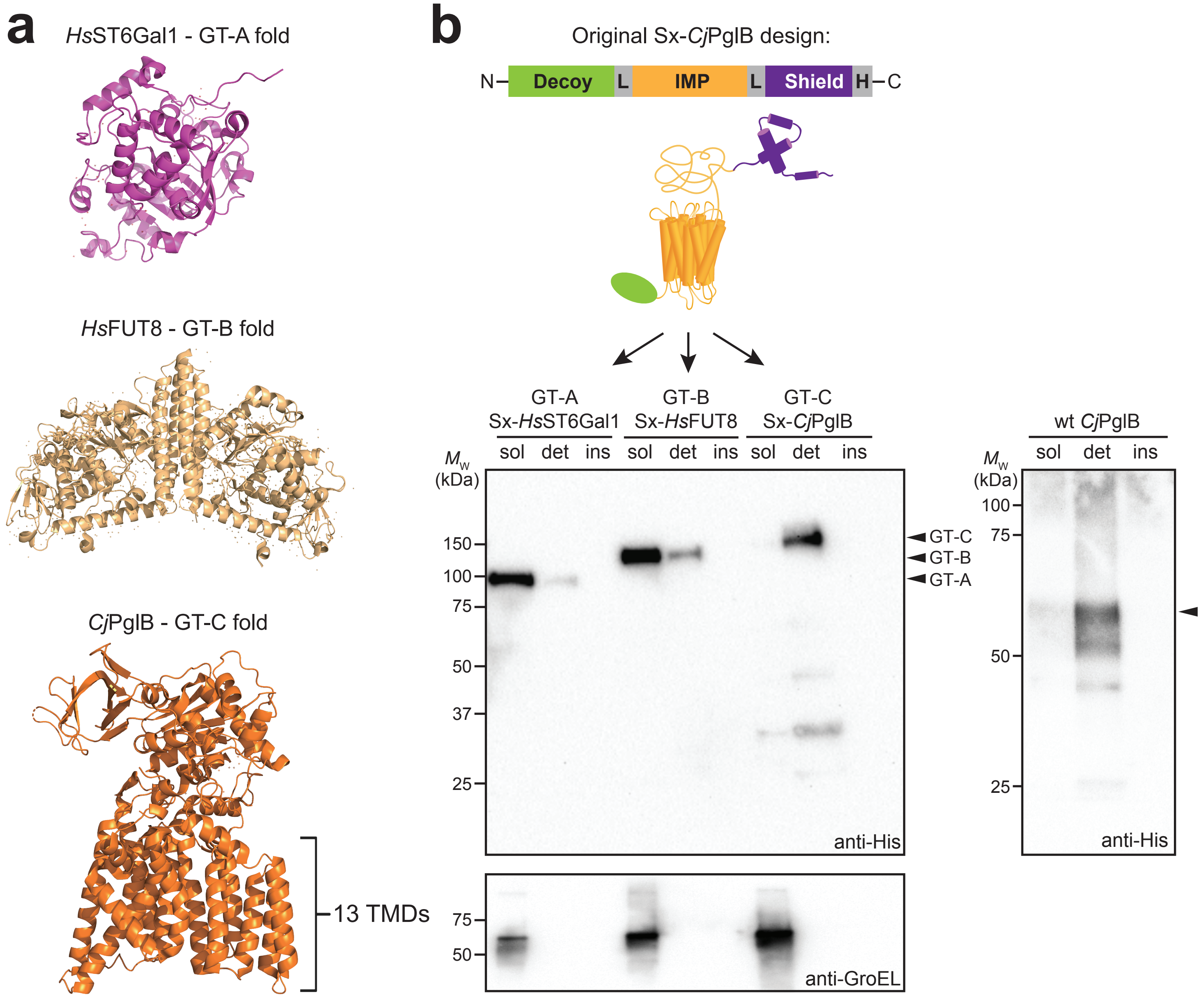


**Supplementary Figure 1. SIMPLEx-reformatted expression of different GT superfamily enzymes.** (a) Crystal structures of representative GT-A, GT-B and GT-C superfamily members (from top to bottom): *Homo sapiens* ST6Gal1 (*Hs*ST6Gal1; PDB 4JS1), *H. sapiens* FUT8 (*Hs*FUT8; PDB 6TKV), and *C. lari* RM2100 PglB (*Cl*PglB; PDB 5OGL). (b) (top) Schematic of original SIMPLEx design with integral membrane protein (IMP) target – GT-A, GT-B or GT-C – expressed as sandwich fusion between N-terminal ΔspMBP domain (decoy) and C-terminal ApoAI* domain (shield). (bottom) Immunoblot analysis of soluble (sol), detergent soluble (det), and insoluble (ins) fractions derived from *E. coli* BL21(DE3) cells expressing SIMPLEx-reformated *Hs*ST6Gal1, *Hs*FUT8 and *Cj*PglB from plasmid pMLBAD. Shown at right is immunoblot of same fractions from BL21(DE3) cells expressing wild-type (wt) *Cj*PglB without any fusion partner. Blots were probed with anti-polyhistidine antibody (anti-His) to detect each of the expressed constructs and anti-GroEL antibody to confirm quality of fractionations and equivalent loading of soluble fractions. Molecular weight ($M_w$) markers are indicated at left. Black arrows indicate protein product bands based on expected $M_w$ of ech construct. Results are representative of three biological replicates.

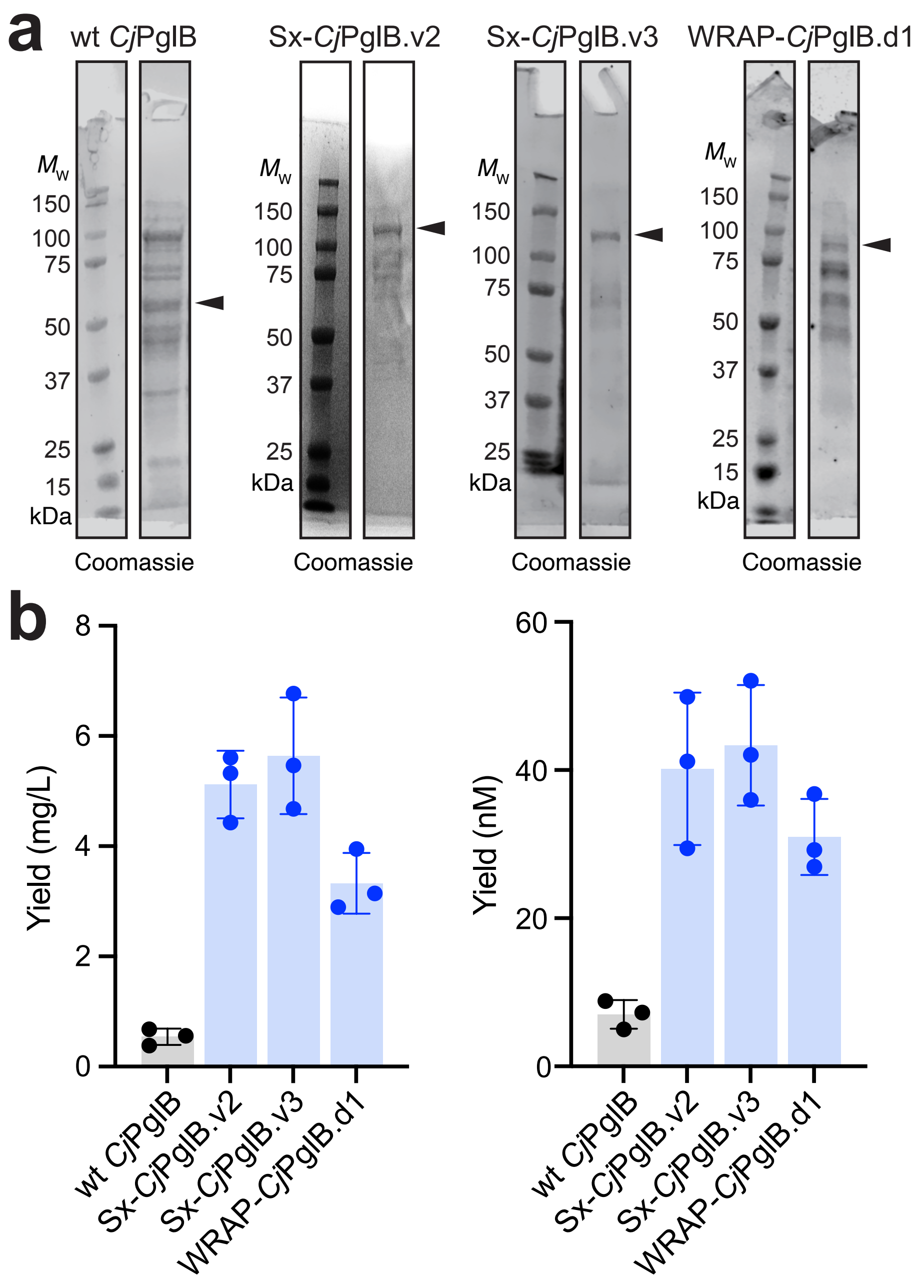


**Supplementary Figure 2. Expression and purification of *Cj*PglB constructs.** (a) Representative Coomassie-stained SDS-PAGE gels of wt *Cj*PglB, Sx-*Cj*PglB.v2, Sx-*Cj*PglB.v3 and WRAP-*Cj*PglB.d1 following expression in 1-L culture of BL21(DE3) cells and purification by $Ni^{2+}$-affinity chromatography. SIMPLExed and WRAPed constructs were purified from total cell lysates while unfused wt *Cj*PglB was purified from the detergent-solubilized membrane fraction. An equivalent volume was loaded in each lane. Results are representative of three biological replicates. Molecular weight ($M_w$) marker is shown on the left. Black arrows denote full-length expression products. For each construct, depicted lanes were from the same gel but with extra lanes removed (b) Soluble protein titers for each construct in (a) determined on a mass basis (mg/L) and a molar basis (nM). Titer values are the mean of biological replicates ($n$ = 3) ± SD.

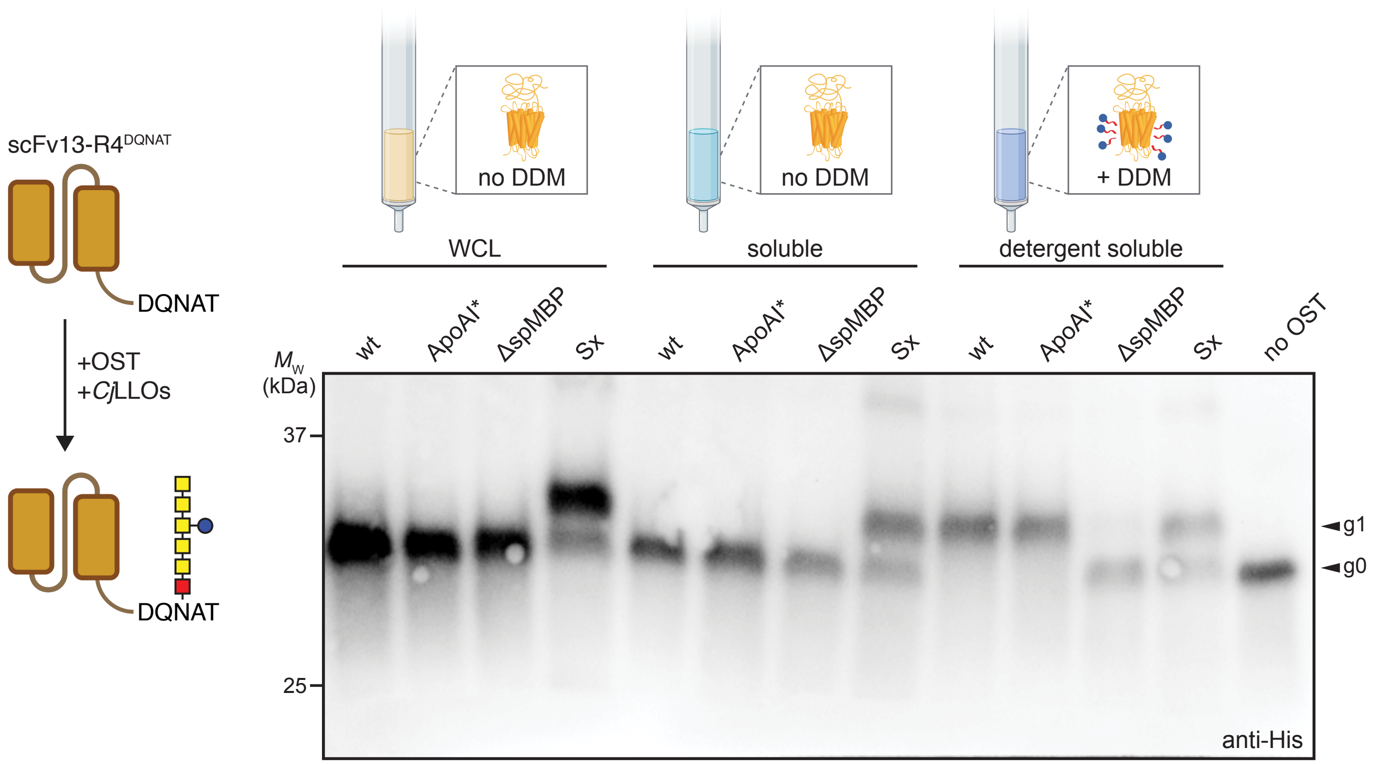


**Supplementary Figure 3. Enzymatic activity of *Cj*PglB constructs derived from different cellular fractions.** Immunoblot analysis of IVG reactions in which different *Cj*PglB constructs were purified from the whole cell lysate (WCL), soluble (sol) or detergent-solubilized membrane (det) fractions of BL21(DE3) cells and incubated with organic solvent-extracted *Cj*LLOs and purified scFv13-R4$^{DQNAT}$ acceptor protein. IVG reactions performed without *Cj*LLOs or *Cj*PglB served as negative controls. Blots were probed with anti-His to detect the acceptor protein. Arrows denote aglycosylated (g0) and singly glycosylated (g1) forms of scFv13-R4$^{DQNAT}$. For all immunoblots, molecular weight ($M_w$) markers are indicated at left and results are representative of biological replicates ($n = 3$).

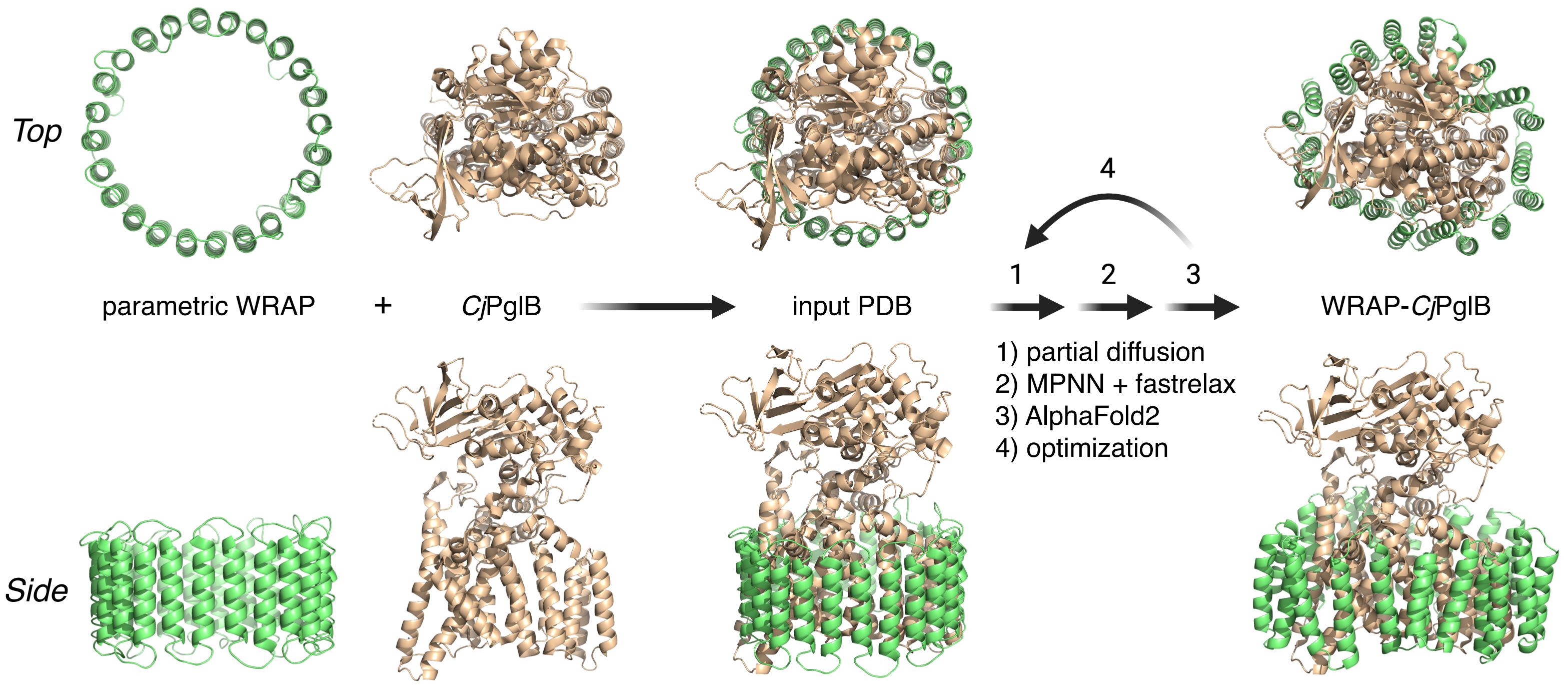


**Supplementary Figure 4. The WRAP design pipeline.** Top and side views of the WRAP design pipeline. The parametric WRAP was generated and positioned around the *Cj*PglB (PDB 5OGL) enzyme's membrane spanning region using sushimaki, and this composite structure was used as the input for partial diffusion. The WRAP sequence and backbone geometry were iteratively optimized (4) while the *Cj*PglB backbone and sequence were held fixed using cycles of partial diffusion for refining the WRAP backbone (1), SolubleMPNN plus Rosetta FastRelax for sequence design (2), and AlphaFold2 initial guess structure prediction for design validation (3).

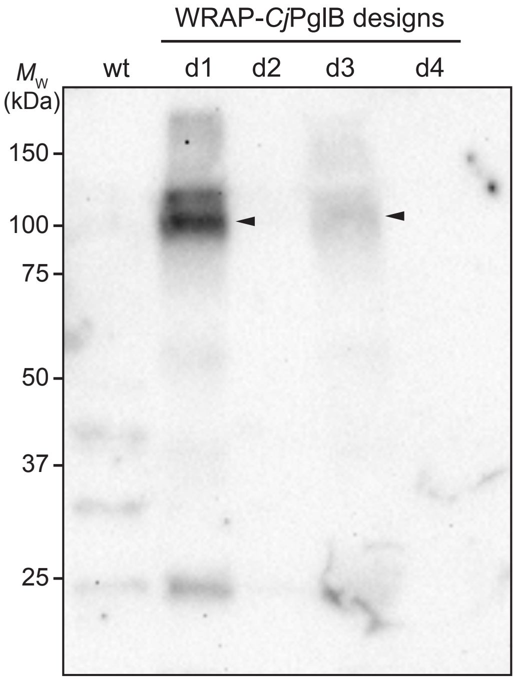


**Supplementary Figure 5. Evaluation of WRAP designs for solubilizing *Cj*PglB.** Immunoblot analysis of soluble fractions prepared from BL21(DE3) cells expressing each of the indicated WRAP-*Cj*PglB constructs. Unfused wt *Cj*PglB prepared identically was included as a control. An equivalent amount of total protein was loaded in each lane. Blots were probed with anti-His to detect each of the expressed constructs Black arrows denote full-length expression products. Molecular weight ($M_w$) markers are indicated at left and results are representative of biological replicates ($n = 3$).

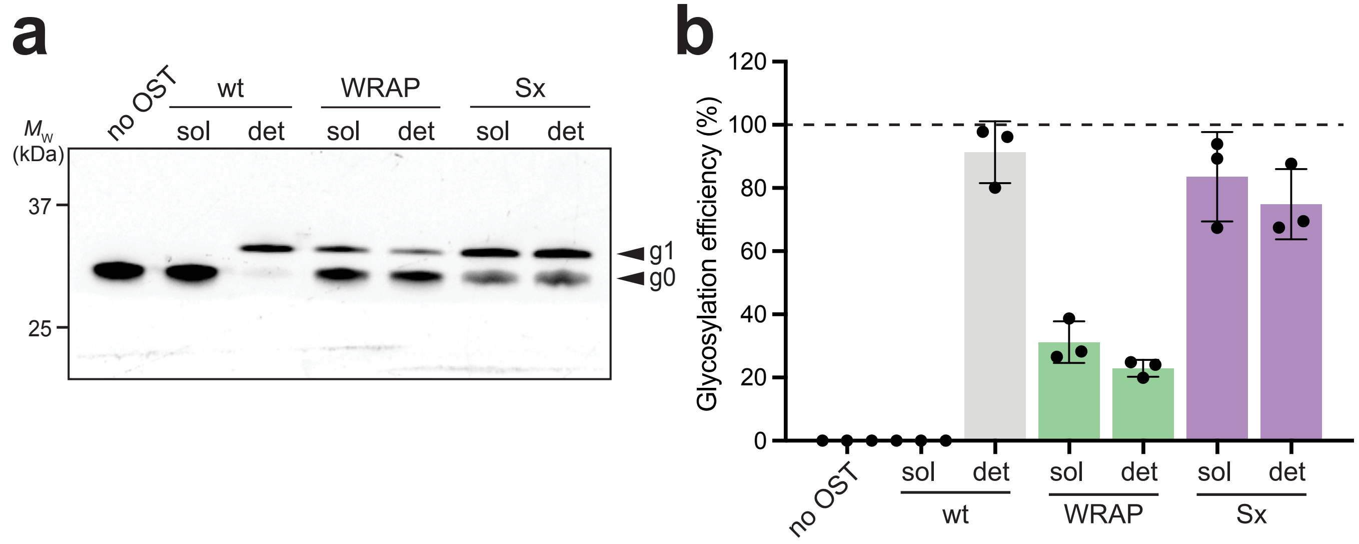


**Supplementary Figure 6. *In vitro* glycosylation of TAMRA-labeled peptides by water-soluble OSTs.** (a) Tricine SDS-PAGE analysis of IVG reactions in which 0.5 µM of TAMRA-labeled peptides and organic solvent-extracted *Cj*LLOs were incubated with ~1 µM of either wt *Cj*PglB, WRAP-*Cj*PglB.d1 or Sx-*Cj*PglB.v3. Each *Cj*PglB construct was purified from soluble (sol) or detergent soluble (det) fractions of BL21(DE3) cells. IVG reactions in which no OST was added served as a negative control. Black arrows denote aglycosylated (g0) and singly glycosylated (g1) forms of scFv13-R4DQNAT. Molecular weight ($M_w$) marker is indicated at left and results are representative of biological replicates ($n$ = 3). (b) Glycosylation efficiency was determined based on densitometric quantification of the fraction of peptide glycosylated expressed as the ration g1/[g0+g1]. Data are mean of biological replicates ($n$ = 3) ± SD.

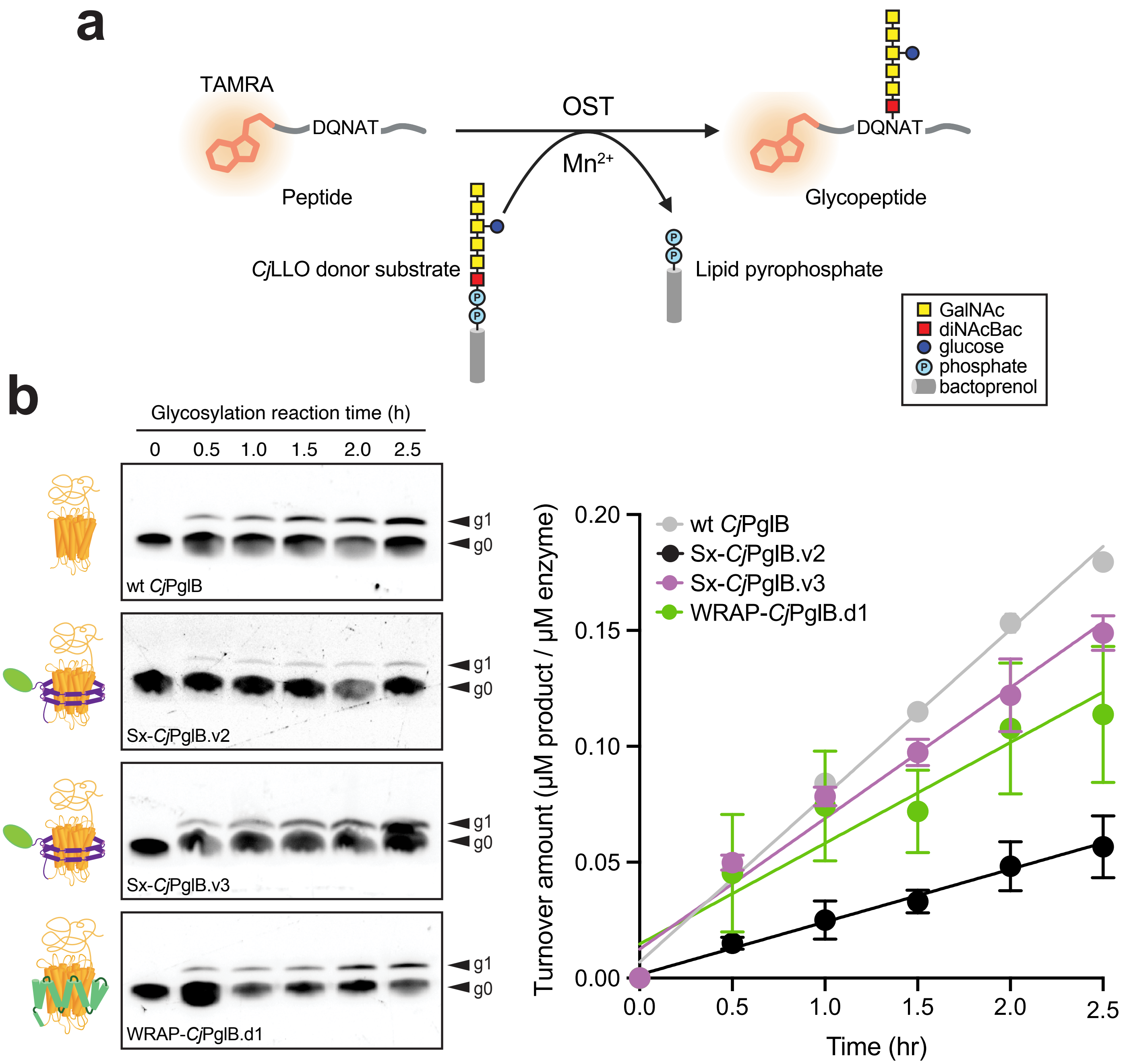


**Supplementary Figure 7.** ***In vitro* glycosylation of TAMRA-labeled peptides by water-soluble OSTs.** (a) Schematic of TAMRA peptide IVG assay. Incubation of TAMRA-labeled peptides with OSTs, CjLLOs, and divalent metal ions (e.g., Mn2+) results in transfer of heptasaccharide glycans to form glycopeptide products. (b) Tricine SDS-PAGE analysis of IVG reactions determined by quantification of TAMRA-labeled peptide substrate and product over time using 0.5 µM peptide substrate and ~1 µM of each OST. Graph at right depicts turnover amount (µM of glycosylated peptide per µM of *Cj*PglB construct) versus time using IVG reaction data. Data fitting was by linear regression, and turnover rates were calculated from slope using Prism 10 for MacOS (version 10.3.0). Tricine SDS-PAGE gels in panel (b) are representative of three biological replicates. Data in corresponding graphs are average of biological replicates ($n$ = 3) ± SD.

**Supplementary Table 1.** Top 5 WRAP sequences selected for experimental testing.

| | |
|---|---|
| 5OGL_5240010 | KKLEELLIKTLLSALKDVSKYKEYAEEVLKYISELLKDPLVVEIVKTFIEAM<br>VAEVQGEKDKDKIIELLKKTTDLLAELAEDEENKNLILEVTETIIKILKMSK<br>EENEKALLLLAKTLIKLLAKLIGDEAAMMAAVAFMFVSSPELREELLETA<br>KEILDEESYLAVKSLGELLILAEKGDKEGLFEKFKEFMEEAKKFDKYKYL<br>GLSLLALLIVATSLDRELALKAFEYFLEVIKPLLDEESYKLISEAIEILKEEK<br>DDKKAMEKLAELMIEYLIELNEDEEVKKVVKEAVEVAKKGDEEAEKEAI<br>AKLLLKAAKIVSKVTGDEEILELTKRLIEQLKELGLPLPGFVFLGLLGLALI<br>VFGPIGKGIAKAFVVLGEKRGESIEEIIKALVELIKEAENPMEQAGIMIVL<br>MFIAVLLTGDGKMEEMKFAVELLIEVAEDEIVKLTMKYFGEALENIKDAE<br>KVMEITVKLLVETTLKII |
| 5OGL_522001 | NELVELLIEYFLAAIKDPSKAKEYAEKIIKYIEEIIKNPEVSEIVRLYVEAMV<br>AELQGEKDREKLIELIKKFTNALANLAEDEEVKNTILEVTETVLKILTMSK<br>EQADEAVLLLVETLIRLLNRLIGSEAAAMAAVAFAFVASPELQERILEVA<br>REVLNEEEYKFVETLVRLIILLQRGDYEGAFEVFKEFLEWAKSWDKYKF<br>LWLSLLAALIALTGLPRELALKAAKYFIETIKDLLDEESLNLIEKALKILEEK<br>KDEKEAMLALAKLMIEYLIELNEDEEVKEVVKKAVEVTEEGDVEKEKEA<br>IAELLLKAAEIVSKVTGDKEIVEVMKKFVEALLELGFPLPGFAFLGLLGLA<br>LLVFGKIGFGMAKALVVLAEQRGKSMEEIIDALVELAKEAKDPLEQAGI<br>MIVLIGLAVLALGDGKKDLTEYAMEKLIEIAKDEIVKLTMKYFKEIMENIK<br>DAEKVMELTIKLLVETTFKLF |
| 5OGL_412003 | KEEIELITEAIIELVLNPEKYEEVIKKLKELIKKIVPDELVQKFLQSYVEAVL<br>LEASGNKDLEKLKELIEKAAKYLSELAEDEKVKKLIEEVTKTVLELLEMK<br>KEESEKALTKLFVTFIKLINLMNDEDSTFLAAVAFGFVADPETAEAIVEV<br>AKKLLDKKKAEVLEYLIKIWIYLQKGDYEKALEYFEKAIEIAKEFPRDEAL<br>WLILLIALLTLTGMPEELGLKIVEKVLEIFKELSDEEFVNLVEKAIEILKES<br>KDKEEAMYKLAEFMIEELSKLSEDEEVVEIVKEALEVTKEKDEEKLLETT<br>KKLLEEMARIAAKITGDEEFLEEMKKLIEYLVELGMSVEGLAFLSLFALA<br>LYLTSKFGYGLIKAFNVLYENRDKGIKAQVEALVKLYKEEEDELVRAAIM<br>MIIIGLAVLAALEGDWETAEYAMKLLEEAAEDEIERLVIRYVAEILKAIGD<br>GEKVLALTIRFAIEATYKAI |
| 5OGL_522007 | NELVNLLIEYLKAAIEDPEKAIEYAEKIIKLIKEIVKNPEVSKIVELYIKALTA<br>ETLGEKDKEKLIKLIKEFTNALANLAEDEEVKNLILRVTETVLKILKMEKE<br>QADEALLLLVETLIRLLAELIGDEAAMMAAVAFMFVASPELQERILEVAK<br>EVLNEEEYKAVETLTKLIILAQKGDYEGALEVFKEFLEWVKSLDKYKAW<br>WLSLLAFLIVATSLPRELALEAAEYLIETFKDILDEESLKLLEEALEILKKV<br>KDPKEAMLALAELMAEYLMKLNEDEEVNEVVRKAVEVTRKGDVEAEK<br>EALAELFLKAAEIVSKVTGDEEIVEVMRRLLRLLRELGLPLPGFAFLSAL<br>GLALLVFGPIGFGMAKALVVLGETRGKSIEEMIDALVALAKEEKDPLKQ<br>AGIILVMIGVAVLALGDGKKELTDYAMDKLIEVAEDEIVKYTMEKFKEIM<br>KNINDAKKVMELTIELLVKSTFKIF |
| 5OGL_614004 | YEFVARLLLMIVLLALGKADEALEMAIKILKALAEAGHPAAEEFKKMAEV<br>FEKVKSESINIKRLAVALFLLGLIAGDTKGMLNTIVELIELLLNKEKLEEIA<br>KLALEYAAEEYRPLVEKLIELILKLRNKSKEEIEKLVKEAVEIALKLADPET<br>QERIKRLYEAYKEGIEAIITTVIEELIEEVAKIPELSELATYFLEKLREALEK<br>GDYEKMLLVYVAFLLALGLVDLISVERIKEILDEVLKEKDAVDGLVKLVK<br>LFGLAIMEAAGLGELFKKAIELVEKGDLAGAFILILEEMAKKYTPEMSKM<br>AELLLEALKKALEGASSEELEKLLKELKEIASEVNSKTLVDLAYDLILAAF<br>TGDEEKILEIIEKKIKELGETEAAKILLEFVKLFIEIN |
